# Prediction error drives associative olfactory learning and conditioned behavior in a spiking model of *Drosophila larva*

**DOI:** 10.1101/2022.12.21.521372

**Authors:** Anna-Maria Jürgensen, Panagiotis Sakagiannis, Michael Schleyer, Bertram Gerber, Martin Paul Nawrot

## Abstract

Predicting reinforcement from the presence of environmental clues is an essential component of guiding goal-directed behavior. In insect brains, the mushroom body is central to learning the necessary associations between sensory signals and reinforcement. We propose a biologically realistic spiking network model of the Drosophila larva olfactory pathway for the association of odors and reinforcement to bias behavior towards approach or avoidance. We demonstrate that prediction error coding through the integration of currently present and expected reinforcement in dopaminergic neurons can serve as a driving force in learning that can, combined with a synaptic homeostasis mechanism, account for experimentally observed features of acquisition and loss of associations in the larva that depend on the intensity of odor and reinforcement and temporal features of their pairing. To allow direct comparisons of our simulations with behavioral data [1], we model learning-induced plasticity over the complete time course of behavioral experiments and simulate the locomotion of individual larvae towards or away from odor sources in a virtual environment.

## Introduction

Goal-directed behavior in dynamic situations benefits from the ability to predict future conditions in the environment from the occurrence of sensory clues. In insects, the mushroom body (MB) is the central brain structure for multi-sensory integration, involved in memory formation and recall [2, 3]. It is at the core of learning and retaining valuable associations between sensory inputs and reinforcement in the synapses between the MB intrinsic and its output neurons [4–7].

One of the underlying mechanisms is associative learning, a process that gradually establishes a relationship between two previously unrelated elements. In classical conditioning, the conditioned sensory stimulus (CS) obtains behavioral relevance through its concurrence with the reinforcing unconditioned stimulus (US), an acquisition process depending dynamically on their spatiotemporal proximity. The temporal evolution of this process has been formalized in the Rescorla-Wagner (RW) model (eqn. 1) [8].

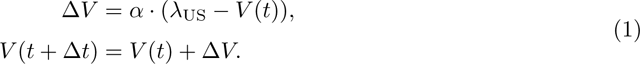

Here, a CS obtains predictive power of concurrent or successive US [8], that depends on the strength of the already acquired association between the CS and US *V* (*t*), allowing for anticipatory behavior to the expected US [9, 10]. The acquisition of this association terminates when the US is fully predicted. Until then, the change in associative strength Δ*V* is proportional to the difference between the maximum associative strength (or asymptote) *λ*_US_ and the current associative strength *V* (*t*) (eqn. 1). The maximum associative strength is a property of the US, determined mainly by the intensity of the reinforcement. While the current associative strength *V* (*t*) is defined by the shared learning history of CS and US [8]. The concept of prediction error (PE) [11] is a derivative of the Rescorla-Wagner model [8]. The error signal equals the difference between the current *λ*_US_ and the predicted value of US *V* (*t*). Over the course of the memory acquisition/training phase, the pace of learning, which can be formalized as the slope of the acquisition curve, decreases as the PE is reduced, minimizing the driving force for changes of the association [8, 11]. This difference is multiplied with a learning rate parameter (*α*), here combined for the CS and the US (eqn. 1).

This continuous optimization of predictions, guided by the PE, could allow animals to efficiently adapt their goal-directed behavior in dynamic environments. Among the most relevant associations to be learned are those that enable the prediction of reward or punishment. Dopaminergic neurons (DANs) have long been known to encode information about reward and punishment. These types of neurons respond to the presence of rewards and punishments in the environment, both in vertebrates [12–17], as well as invertebrates [18–21]. The electrical stimulation or optogenetic activation of DANs induces approach or avoidance both in vertebrates [22–25] and invertebrates [20, 26–32]. In adult [5, 6, 33, 34] and larval [20, 35, 36] *Drosophila* this approach or avoidance learning is facilitated by the modulation of MB output synapses by DAN activity. Ultimately DANs do not only signal the presence of rewards or punishments but have also been suggested to encode PE in various vertebrate species [16, 37–40] and might have a similar function in insects [19, 20, 32, 34, 41–44].

We utilize our spiking model of the *Drosophila* larva MB in one brain hemisphere that forms associations of odors with reinforcement to further test the hypothesis that PE coding within this circuit takes place in DANs that receive input from the output neurons of the MB or their down-stream partners [20, 45–47], that might provide feedback to the DANs. Beyond the scope of similar models [20, 48–51](see Discussion, section: Comparison with other MB models), we demonstrate that this mechanism can reproduce the experimentally observed findings on the acquisition of associations of odors with reinforcement in a time-resolved manner [1]. To facilitate direct qualitative and quantitative comparisons with animal behavioral data, we couple this model with a realistic locomotory model of the larva [52] that captures the effects of learned associations on chemotactic behavior in individual animals.

## Results

### Connectome-based circuit model of the larval olfactory pathway

The network architecture of our model (fig. 1 A) is based on the anatomy of the olfactory pathway in one *Drosophila* larva brain hemisphere [20, 29, 53, 54] (for more details see Methods, section: Network model). Peripheral processing is carried out by 21 olfactory receptor neurons (ORNs), each expressing a different olfactory receptor type [53, 55, 56]. ORNs form one-to-one excitatory synaptic connections with 21 projection neurons (PNs) and 21 local interneurons (LNs) in the antennal lobe [53]. Each LN connects with all PNs via inhibitory GABAergic synapses, establishing a motif for lateral inhibition within the antennal lobe. The 72 mature larval Kenyon cells (KCs) [54] are the excitatory principal cells of the MB. Each KC receives excitatory input from 2-6 randomly selected PNs [54]. The KCs are subjected to feedback inhibition, provided via the GABAergic anterior paired lateral (APL) neuron, which receives input from all KCs [29]. Only mature KCs, characterized by a fully developed dendrite, are included in this model, yielding a complete convergent synaptic KC *>*APL connectivity. The output region of the MB is organized in compartments, in which the KC axons converge with the dendrites of one or few MB output neurons (MBONs) [20, 54]. Our model assumes two MBONs from two different compartments that are representative of two different categories of output neurons of the MB that mediate either approach or avoidance [4–6, 33, 35, 36, 57–59] with a single MBON each. Both MBONs receive excitatory input from all of the KCs to fully capture the information that is normally represented by the complete set of MBONs. Each compartment is also innervated by a single DAN, signaling either reward or punishment and targeting the KC*>*MBON synapses to facilitate learning (for a discussion of all simplifications compared to the animal brain, see Methods, section: Network model).

**Figure 1:**
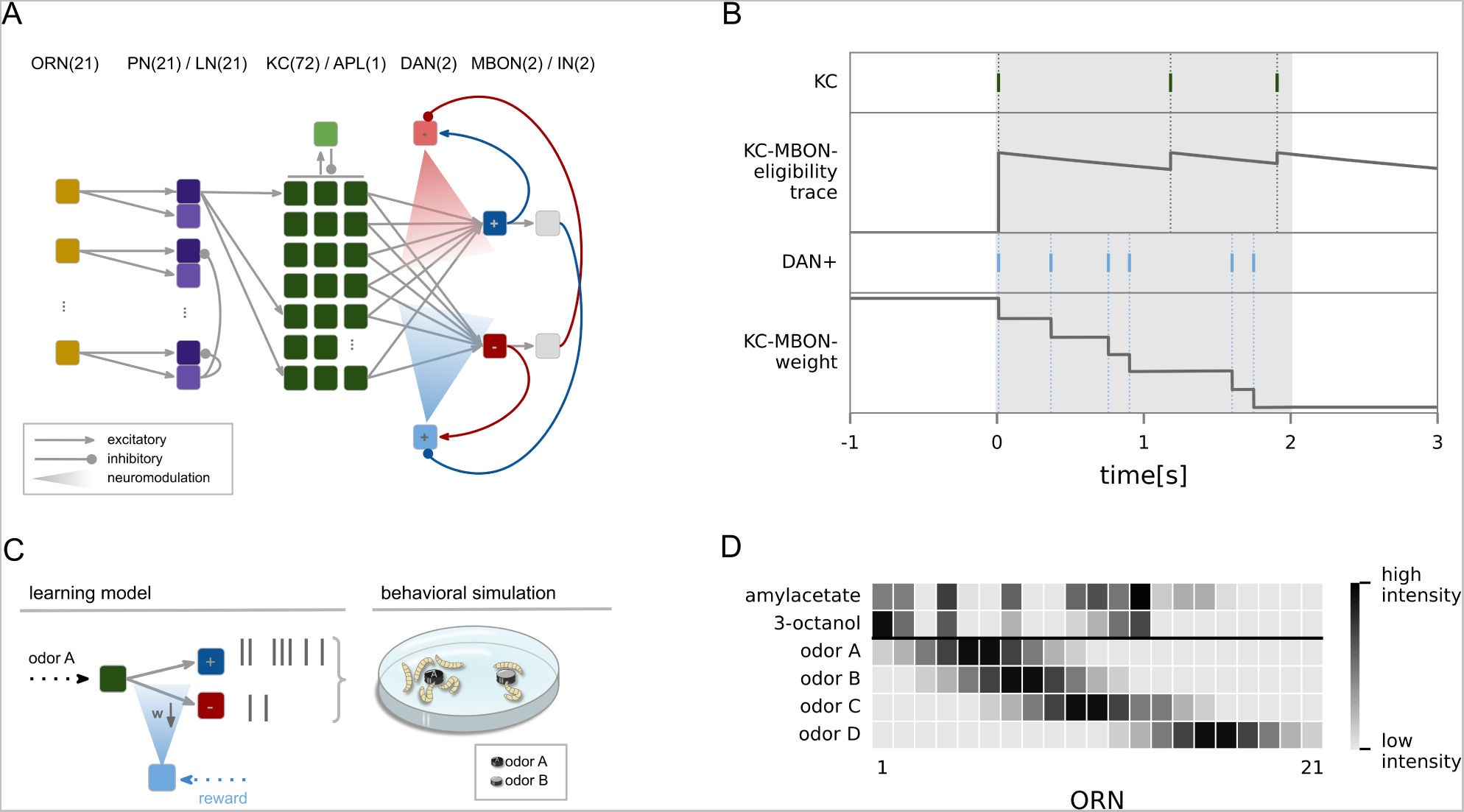
Network mechanisms. (A) Network model of the *Drosophila* larva olfactory pathway including all neurons and connections implemented. One-to-one feed-forward connections between 21 olfactory receptor neurons (ORN) and 21 projection neurons (PN)/local interneurons (LN) and from 2-6 PN to each of the 72 Kenyon cells (KC). Lateral inhibition from each LN innervates all PNs and recurrent feedback inhibition from the anterior paired lateral (APL) neuron is provided onto all KCs. The MB output region is organized in two distinct compartments. The upper compartment holds the approach encoding MBON_+_ and is innervated by the punishment mediating DAN_-_, the lower compartment holds the avoidance mediating MBON_-_ and is innervated by the reward mediating DAN_+_. Each DAN can exert a neuromodulatory effect on the plastic KC*>*MBON synapses within its compartment. MBONs provide excitatory and inhibitory (via gray interneurons) feedback to the DANs. (B) Sketch of synaptic weight change at a single KC*>*MBON synapse with respect to the synaptic eligibility trace elicited by KC spikes and the occurrence of reward-triggered spikes in DAN_+_. Amylacetate is paired with a reward for 2s (gray shaded area). (C) To generate simulated larval behavior in the petri dish during the test phase of the learning experiments, we utilized our locomotory model [52], based on the behavioral bias (eqn. 4) acquired by the MB model during the training phase. The behavioral bias is used directly as input to the locomotory model. (D) All odors (see Methods, section: Sensory input) were used in the experiments. Naturalistic odor patterns for amylacetate and 3-octanol as well as four artificial patterns (odorA,odorB,odorC,odorD) with varying distances (see Methods, section: Sensory input) from odorA. Each odor activates a different set of input neurons with a different spike rate, as indicated by the color bar.

### Learning through KC*>*MBON plasticity

We assume that the KC*>*MBON synapses undergo plasticity, based on strong experimental evidence in larval [20, 35, 36] and adult flies [5, 6, 34]. This plasticity requires the convergence of the sensory pathways in the form of KC activation and of the reinforcing pathway, mediated by neuromodulatory DAN signaling at the synaptic site. We employ a two-factor learning rule (eqn. 2) at each KC*>*MBON synapse (fig. 1 A,B). The first factor is expressed in the pre-synaptic KC activation by an odor, tagging the synapse eligible for modification. This is modeled via an exponentially decaying eligibility trace *e_i_*(*t*), which is set to a 1 whenever the respective KC elicits a spike (fig. 1 B). The decay time constant determines the window of opportunity for synaptic change. The presence of reinforcement (reward or punishment) constitutes the second factor and is signaled by the reward-mediating DAN_+_ or punishment-mediating DAN_-_, respectively. Spiking of the DAN provides a neuromodulatory reinforcement signal *R*(*t*) to the synaptic site. If a DAN spike coincides with positive eligibility at the synapse, the respective synaptic weight is reduced. At each synapse *i*, the reduction of synaptic weight Δ*w_i_* depends on the learning rate *a* (table S1) and is proportional to the amplitude of the eligibility trace *e_i_*(*t*) (fig. 1 B):

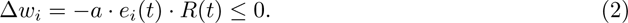

We introduce a synaptic homeostasis mechanism (eqn. 3) that modulates the effects of plasticity at each KC*>*MBON synapse to account for the experimentally observed loss of a learned association when reinforcement is omitted [41, 60, 61] and to ensure continued input to both MBONs. With each MBON spike, the current weight *w_i_* of each respective KC*>*MBON synapse is increased, proportionally to the extent to which the weight differs from its original value *w_init_* (table S1) and multiplied with a homeostatic regulation factor *h* (table S1). This mechanism serves as an implementation for the loss of the association when the reinforcement is omitted. While reinforcement is present, the learning curve will either continue to rise or remain at the asymptote if already saturated. The interaction of the two mechanisms of learning and unlearning at the level of the individual KC*>*MBON synapses allows to include the loss of learned associations, when continued reinforcement is omitted (see Discussion, section: A mechanistic implementation of the RW model) and also ensures continued input to the MBONs, despite the reduction of input weights over the course of the learning process (eqn. 2). The homeostatic factor *h* hereby serves as an implementation of a time constant of this exponential process. The interaction of synaptic plasticity and homeostatic regulation defines the magnitude of the weight at the next simulation timestep *t* + Δ*t* as

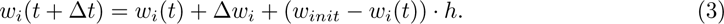

It has been shown in behavioral experiments that specific MBONs encode a behavioral tendency to either approach or avoid a currently perceived stimulus, depending on the acquired stimulus valence [4–6, 36, 57, 58]. In the naive state of our model, all KC*>*MBON synapses have the same initial weights *w_init_* (table S1), and hence the spiking activity of both MBONs is highly similar. Learning alters the KC*>*MBON synaptic weights and thus skews the previously balanced MBON output. This acquired imbalance between MBON outputs biases behavior towards the approach or avoidance of the conditioned odor. To quantify the effect of learning, we compute the behavioral bias BB (eqn. 4) from the firing rates of both MBONs over *T* = 1*s* as follows:

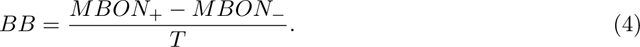

### Implementation of prediction error coding in the KC-MBON-DAN motif

In the larva, many DANs and other modulatory neurons receive excitatory or inhibitory input from different MBONs, either in a direct manner or via one or two interneurons [20]. Based on this observation, we constructed our hypothetical feedback motif (for similar models see discussion section: Comparison with other MB models). In the model, DANs are activated by external reward/punishment signals and also receive excitatory and indirect inhibitory feedback from both MBONs (fig. 1 A). As the initial balance between the two MBON outputs shifts over the course of the training process, the amount of excitatory and inhibitory feedback that DANs receive continues to diverge, allowing the DANs to access the model’s learning history. Ultimately DAN activation signals the difference between the current external activation and the expected activation based on prior learning, implemented as the difference between excitatory and inhibitory MBON*>*DAN feedback. Including this feedback leads to learning curves that saturate when the reward is fully predicted, and the prediction error approaches zero (fig. 2 A,D). This effect disappears, when the feedback circuit is disabled (fig. 2 A). In this case the behavioral bias quickly reaches the maximum value of the measure when the MBON_-_ elicits no more spikes and can not encode further learning. Increasing reward intensity, learning rate or odor intensity (see Methods, section: Experimental protocols) foster a faster acquisition of the association and increases the maximum strength of the association at the same time (fig. 2 A).

**Figure 2:**
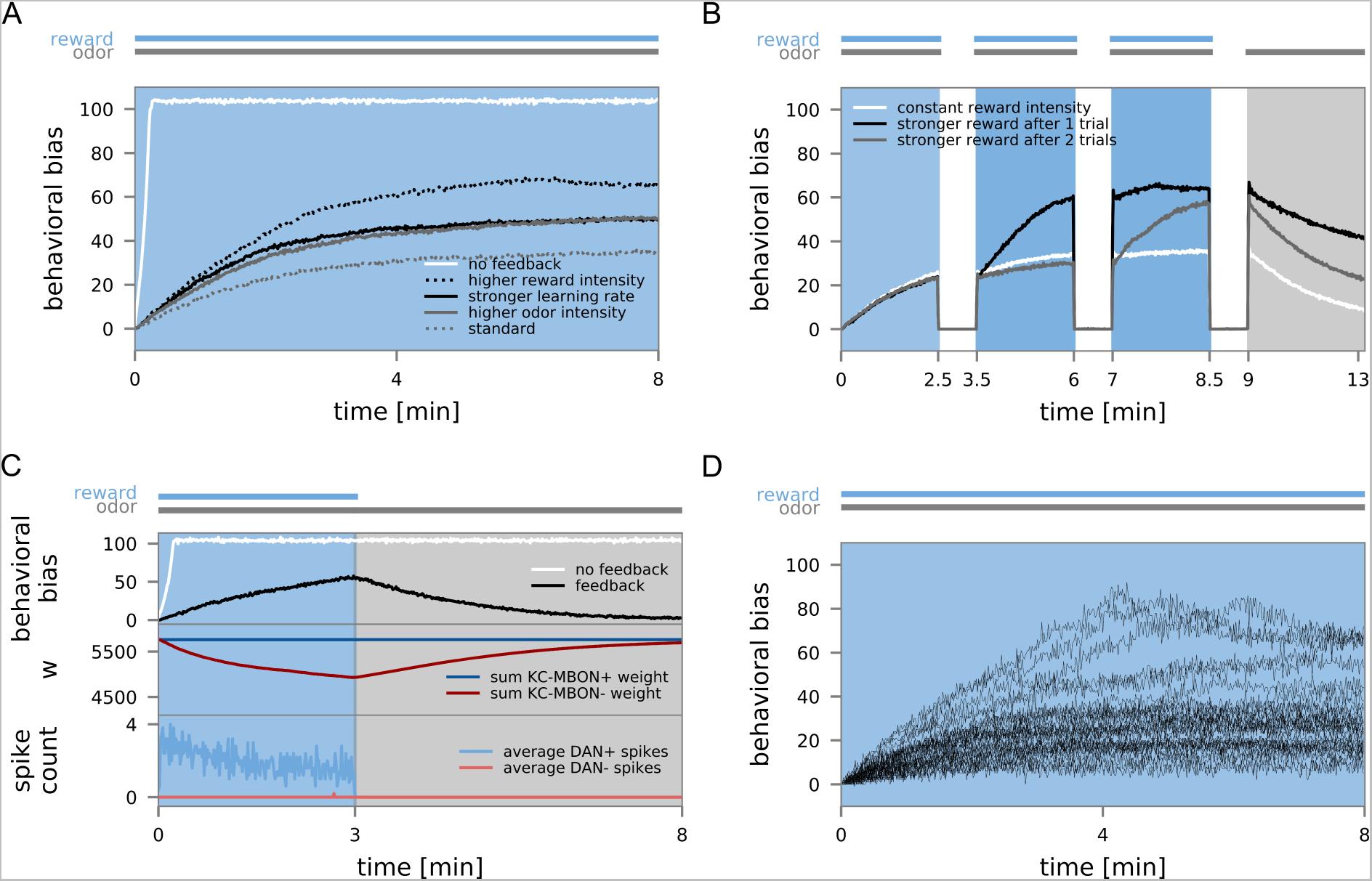
Learning with prediction errors. (A) *N* = 30 model instances were trained with the odor amylacetate (CS) and reward (US, blue background). MBON*>*DAN feedback, the reward/odor intensity, and the learning rate were manipulated in separate experiments. The odor preference (behavioral bias, eqn. 4) was measured continuously in windows of 1 sec and averaged over all model instances. (B) *N* = 30 model instances were trained during three trials with amylacetate and reward (blue background). Reward intensity was either constant across the three training trials (white curve), or enhanced during the third (gray) or the second and third trials (black). The training was followed by a 3 min test phase with odor only (gray background). (C)*N* = 30 model instances were trained with amylacetate and reward (blue background) and then underwent an extended test phase (gray background). (D) Individual acquisition curves for *N* = 30 model instances (standard experiment fig. 2 A).

Increasing the reward intensity after a 2.5 min (black curve), or 5 min (gray curve) of appetitive training, results in a steeper slope of the learning curve and also increases the maximum during training trials of 2.5min duration with increased reward intensity (fig. 2 B). Higher intensity of the reward results in an average DAN spike rate of 39.14Hz(*std* = 1.27(standard deviation)) compared to 33.11Hz(*std* = 1.34).

Additionally, we tested for loss of the acquired association as the reduction in behavioral memory expression, over the course of prolonged exposure to the CS without the US, following initial memory acquisition [8, 62]. We test this in our model experiments by presenting the odor, previously paired with reward, for an extended period of time, in the absence of reinforcement. During the test phase and without the presence of reward to trigger synaptic KC*>*MBON_-_ weight reduction, the extinction mechanism is no longer outweighed by learning and drives each individual weight back towards *w_init_* (fig. 2 C, upper panel). We also demonstrate the interaction of the learning rule with this mechanism in figure S1, where the learning rate remains constant but the magnitude of the homeostatic regulation was manipulated to show that both mechanisms need to be in balance.

### Learned preference and behavior generalize to similar odors

We trained our model by pairing a reward with a single odor for 4min. After the training procedure, we tested the behavioral bias either for the same or a different odor, following the experimental approach used in the larva [63]. Mimicking the experimental data, we show that the odor preference is highest if training odor and testing odor are identical in the case of training with 3-octanol. When amylacetate is used during training, 3-octanol preference is increased (fig. 3 A). Since 3-octanol activates a subset of the ORNs activated by amylacetate (fig. 1 D), some of which with higher rates than in the case of amylacetate, we also tested for generalization using a set of ORN activation patterns with a controlled degree of overlap (see Methods, section: Sensory input, fig. 1 D) and show that with decreasing similarity, the generalization effect to a new odor is diminished (fig. 3 A). Figure 3 B shows the network response to 30sec stimulations with amylacetate and 3-octanol in a single exemplary model instance. On the level of the ORNs, 3-octanol merely activated a subset of the amylacetate-activated neurons. The uniqueness of the odor identities is enhanced in the KC population [64].

**Figure 3:**
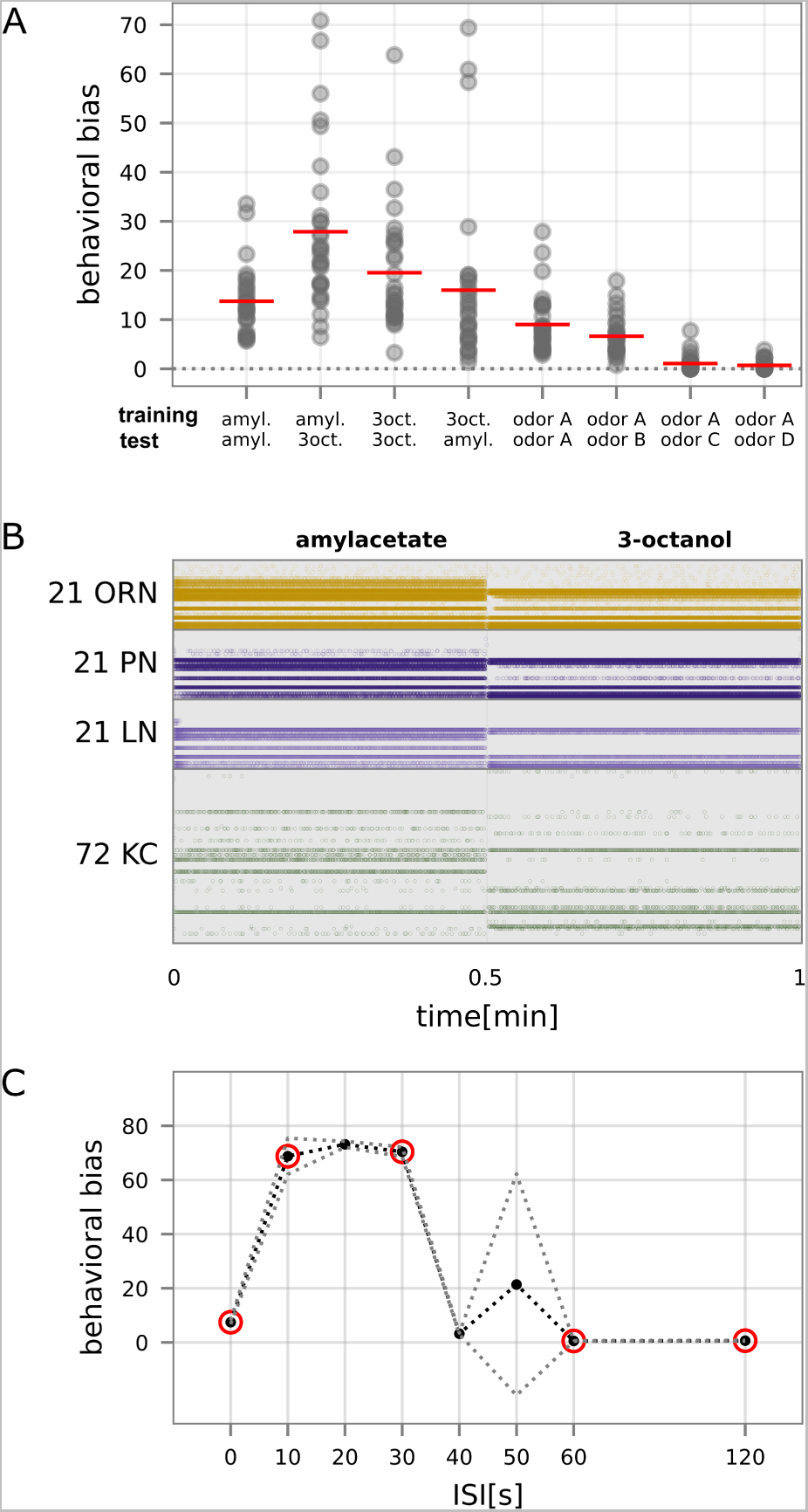
Reward generalization and trace conditioning. (A) The behavioral bias generalizes to odors that differ from the training odor after a 4min training (3min test phase). We conducted simulation experiments with different combinations of training and testing odor, each for 10 groups (gray circles represent the mean of a single group) of *N* = 30 larvae, and red lines indicate the mean between groups. The behavioral bias is highest when the training and the testing odor are the same. (B) Spiking activity in the network during the presentation of amylacetate (left) and 3-octanol (right) in a single naive model instance. (C) Simulated trace conditioning experiments with odor (amylacetate) and reward. Inter-stimulus interval (ISI) indicates the time between odor and reward onset. The black line displays the mean, gray lines the std over *N* = 10 groups of 30 model instances each. Conditions circled in red correspond to the conditions also used in animal experiments [29, 66]

### The model reproduces temporal features in trace conditioning experiments

Including an odor-evoked eligibility trace at the KC*>*MBON synapses allows the model to maintain the sensory odor representation for a time window, during which reinforcement will trigger synaptic change (fig. 1 B). The time window between odor and reward onset (0, 10, 20, 30, 40, 50, 60, 120s) was varied for trace conditioning experiments with a 30s presentation of odor and reward that was repeated three times. A small inter-stimulus-interval (ISI) of 10 to 30s leads to an increase in behavioral bias compared to the complete overlap of odor and reinforcement (fig. 3 C), using the extended window of opportunity for synaptic change triggered by each KC spike. Long ISIs do not lead to learning as the eligibility trace declines back to zero during this time (fig. 3 C). These findings match observations from experiments in larvae [29, 65, 66] with the caveat that the trace in the real larva brain seems to extend for a slightly longer period of time, compared with our experiments.

### The model reproduces paired and unpaired associative conditioning experiments

To test if learning, driven by prediction error, can account for learned larval behavior, we replicated single-trial conditioning experiments performed with larvae [1] in simulation. In these experiments, animals were trained with the odor amylacetate in a single trial of varying duration (1 *−* 8 min). To this end, larvae were placed on a Petri dish coated with an agar-sugar substrate and the odor in two small containers for diffusion in the air (paired training). Either before or following this training protocol, larvae underwent a single trial without sugar and odor. Afterward, the animals were transferred to a new dish with two odor containers placed on different sides (one of them contained amylacetate, and the other one was empty). This paired training was compared with an unpaired protocol with separate (randomized order) presentations of amylacetate and sugar. Following the paired training protocol (odor and reward are presented concurrently), the animals showed a tendency to approach the previously rewarded odor, as measured by the difference in the number of animals on each side at the end of a 3min test phase, divided by the total number of animals. Following the experimental literature, we will refer to this measure as the preference index ([1] eqn. 15). The animals’ preference is relatively consistent across training trials of different duration. Prolonged paired training did not lead to an increase in preference (fig. 5 A). These experiments did not include a test for odor preference before training, but untrained larval odor preference of odors used in learning experiments has been demonstrated elsewhere [67–69]. This paired training was compared with an unpaired protocol with separate (randomized order) presentations of amylacetate and sugar. Here the extent to which animals preferred amylacetate over no odor varied with the duration of the training trial. The longer the duration of the training, the more the preference index decreased from an initially high value but saturated around 2.5min (fig. 5 A).

We aimed to replicate these behavioral experiments on two levels. Firstly, we focused on the direct model output that reflects the strength of the acquired association between amylacetate and reward (behavioral bias, eqn. 4) and later also simulated behavior based on these biases. We simulated both the paired and unpaired training protocol (fig. 4 B). While the unpaired training yielded almost no behavioral bias, the models that underwent the paired training show an increased behavioral bias, that depended on the duration of the training and saturated for longer training duration (fig. 4 B). The simulation results reported in figure 4 B were obtained using odor-naive models that exhibited no odor preference, prior to training. To account for the experimental finding that real larvae often do have an odor preference even without any training [67–69], we readjusted our experiments to include a pre-training period of 10 minutes to start the conditioning experiments with the amylacetate-reward association already established. This adaptation of the protocol leads to results (fig. 4 C) that match the results obtained in real behavioral experiments (fig. 5 A). The paired condition in figure 4 C shows that once the behavioral bias is saturated (fig. 2 A), continued pairing maintains the association, without further increasing it. Unpaired training on the other hand, causes the behavioral bias to decrease and saturate at a lower level. For a discussion of different potential causes of a reinforcement expectation prior to training, please refer to the discussion (Comparison of modeling results to experimental findings). Figure 2 A demonstrates that disabling MBON*>*DAN feedback leads to a learning curve that does not saturate but instead increases with a steep slope until it reaches the maximum value for the behavioral bias eqn. 4) with a MBON_-_ rate of 0. To verify if this PE feedback mechanism is responsible for the difference between maintenance and loss of the association in figure 4 C, we repeated the same experiment with disabled MBON*>*DAN feedback. The behavioral bias overall is much higher, compared to the intact network (fig. 4 B). The maximum is reached before the test phase of even the shortest1min training experiment, with no MBON_-_ spikes elicited.

**Figure 4:**
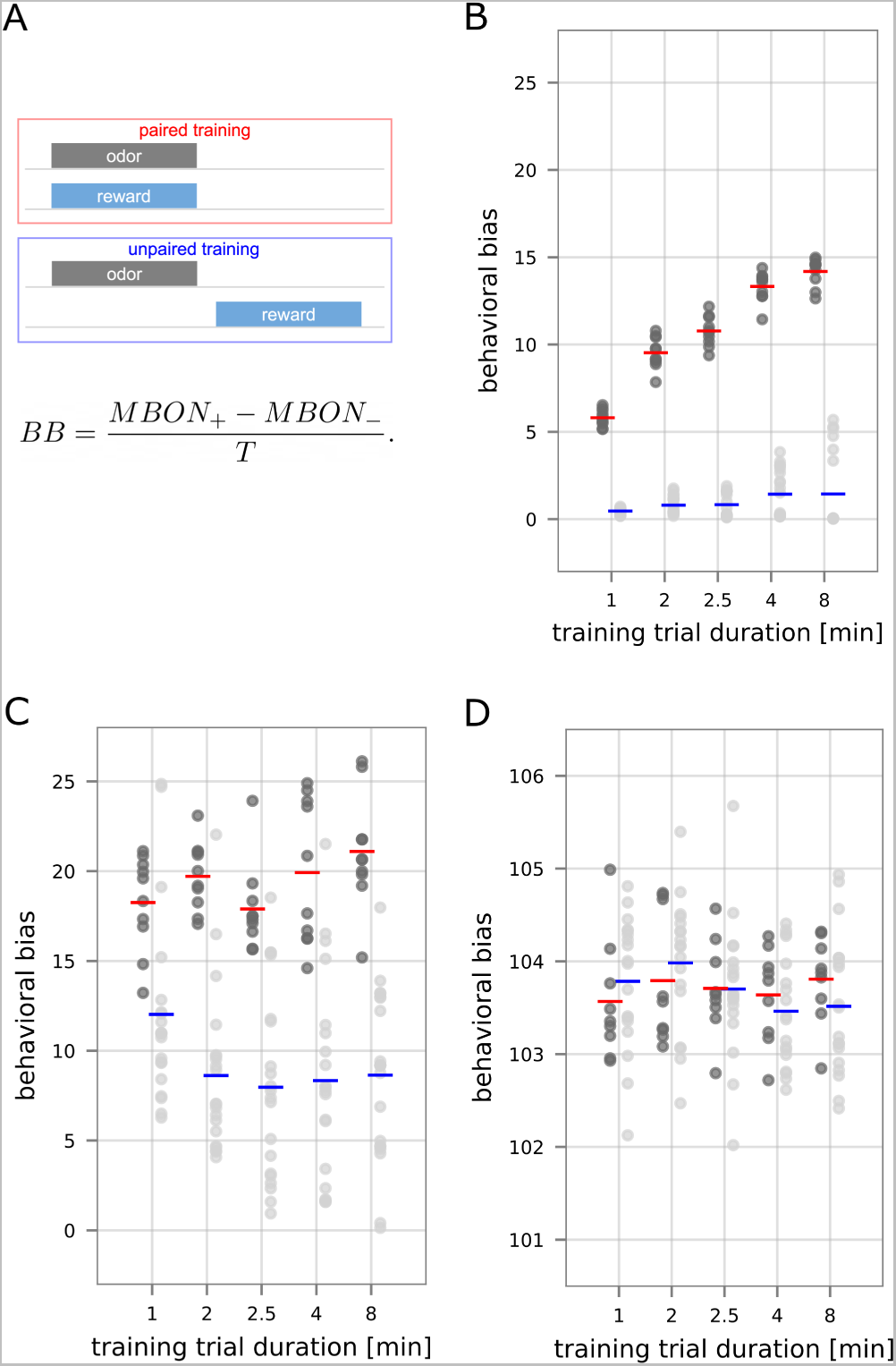
Paired and unpaired learning in the MB model. (A) Schematic overview of the paired vs. unpaired training protocol. (B) The model’s behavioral bias for training with amylacetate and reward for *N* = 10 paired (dark gray, mean in red) and *N* = 10 unpaired (light gray, mean in blue) experiments with groups of 30 modeled larvae each. In the unpaired condition, half of the groups were trained with the odor preceding the reward, for the other half, the reward preceded the odor. (C) Model behavioral bias for amylacetate for *N* = 30 paired and *N* = 30 unpaired experiments with randomized order of odor and reward. Prior to the conditioning experiment the model instances underwent a 10min pre-training period, during which odor and reward were paired. (D) Model behavioral bias for amylacetate for *N* = 30 paired and *N* = 30 unpaired experiments with randomized order of odor and reward. The MBON*>*DAN feedback was disabled. Prior to the conditioning experiment the model instances underwent a 10min pre-training.

Secondly, since the effect of training in lab experiments is quantified behaviorally via spatially defined, group-level metrics (preference index and performance index (eqn. 15,eqn. 16), [1]), we performed behavioral simulations of the testing phase with groups of virtual larvae for both the paired and unpaired condition [1], allowing a straightforward comparison with the animal experiments (fig. 5 A). To this end, we utilized a realistic model for the simulation of larval locomotion and chemotactic behavior [52] that uses the behavioral bias at the output of the MB model as a constant gain factor to modulate the locomotory behavior of individual larvae towards or away from a spatially placed odor source in a virtual arena (see Methods, section: Realistic modeling of larval locomotion). The resulting preference indices, acquired across groups of independently simulated larvae (fig. 5 C), can directly be compared to the experimentally obtained preference indices (fig. 5 A). We also compare performance indices from our simulated experiments (fig. 5 D) with those from the lab experiments (fig. 5 B) and find that the model can replicate these when accounting for the odor preference at the beginning of the experiment.

**Figure 5:**
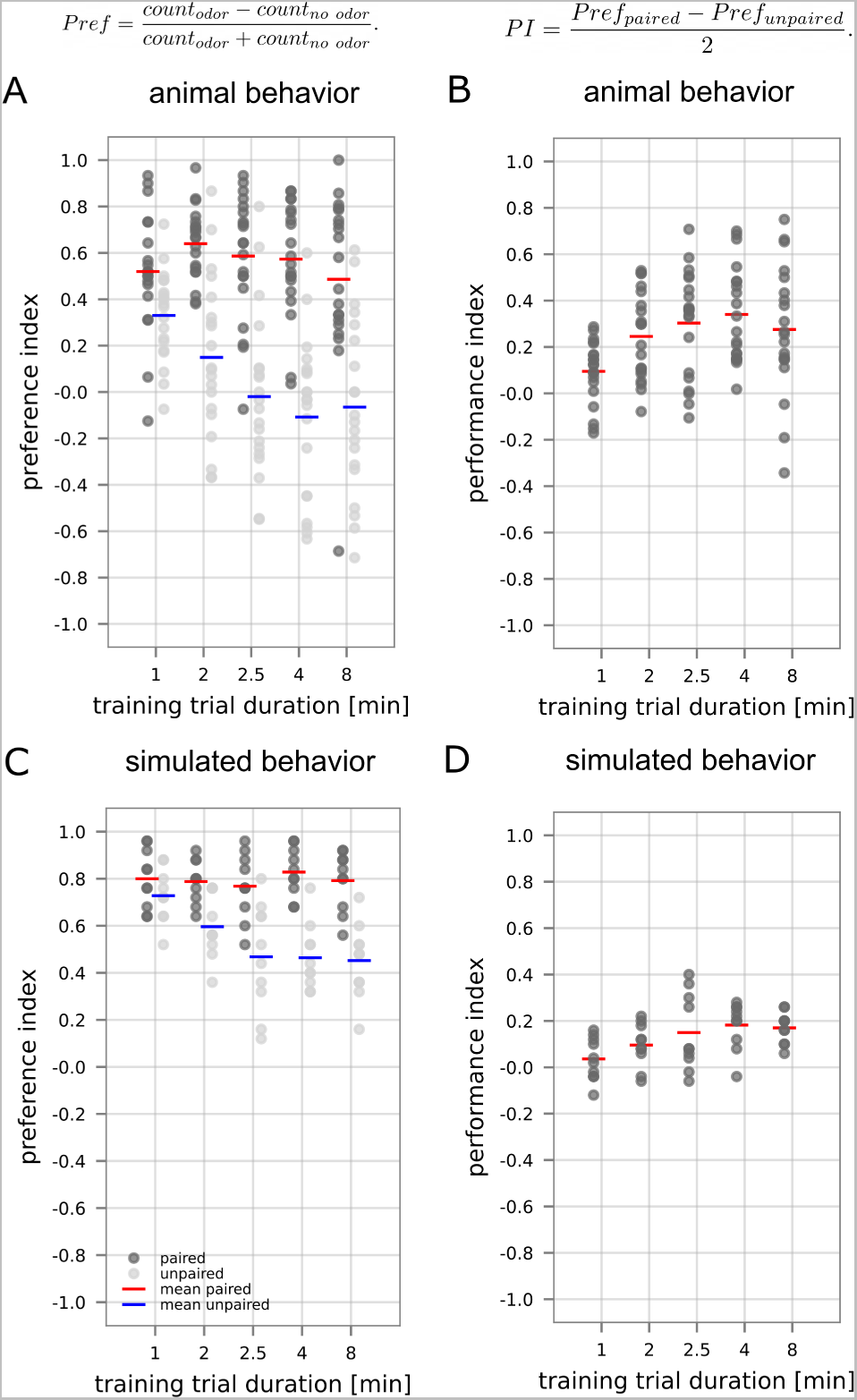
Replicating behavioral experiments with paired and unpaired training. (A) Experimental preference indices for amylacetate for 20 groups of 30 real animals each for paired and unpaired experiments with randomized order of odor and reward [1]. (B) Experimental performance indices for amylacetate computed between preference in paired and unpaired real animal experiments [1]. (C) The simulated behavior is based on the protocol in A. Simulated preference indices for amylacetate for *N* = 10 paired and *N* = 10 unpaired experiments with varied order of odor and reward. (D) Simulated performance indices for amylacetate computed between preference in paired and unpaired simulation experiments.

## Discussion

Seeking rewards and avoiding punishments by predicting change in the environment is a major motivator of animal behavior. Sensory clues can acquire the necessary predictive power to guide behavior through classical conditioning, an associative learning process potentially driven by reward/punishment PE [8, 11], as observed in vertebrates [16, 37–40]. To test the biological plausibility of the proposed PE coding motif in the larval MB and test its capacity to explain behavioral data we implemented a spiking network model of the olfactory pathway, coupled with a simulation of locomotory behavior [52]. We demonstrate that our model of PE coding results in saturating group-level and individual learning curves, where the slope and maximum of the learning curve are determined by the intensity of both the reward and the odor signal. Learning is also influenced by the timing of odor and reinforcement and can be extinguished if reinforcement is omitted during the presentation of the sensory clue. After verifying that this circuit motif enables learning as predicted by the PE theory, we show that it can also explain time-resolved larval behavior in conditioning experiments.

### A mechanistic implementation of the RW model

A number of predictions can be derived from the phenomenological RW model [8] and tested in our mechanistic model thereof. We found that regardless of odor/reward intensity or the model’s learning rate, the strength of the odor-reward association (quantified as the behavioral bias) saturates over time (fig. 2 A), as the strength of the already acquired association *V* (*t*) approaches the maximum value supported by the given reinforcement input (*λ*_US_) (eqn. 1). Consequently, our model’s acquisition curve saturates at a higher value when the intensity of the reinforcement is increased (fig. 2 A,B), as predicted by the RW model, in which a stronger US should result in a higher value of *λ*_US_ [8]. In our model, a higher reinforcement intensity relates to a higher input rate into the respective DAN (see Methods, section: Sensory input) which translates into more frequent DAN spikes within a given window of 1 second, used to compute the behavioral bias (eqn. 4). This defines the asymptote of the learning curve. According to the RW model, increasing either the intensity of the odor or the learning rate *α* [8] should lead to faster acquisition of the association. In our model, the learning rate directly influences the increment of each respective synaptic weight Δ*w*^i^, while an increase of the odor intensity allows for a more frequent execution of the weight update routine, by influencing the eligibility trace (eqn. 2).

The RW model predicts that the omission of reward should result in the loss of the learned association (eqn. 1, [8]). From the equation itself, we can not infer if this loss is due to extinction or forgetting. Extinction, characterized by the possibility of recovery of the association, after its temporary loss [70], has been demonstrated in adult [71, 72] but not larval *Drosophila*. To retain the association for recovery, extinction relies on the formation of parallel memory traces for the acquisition and the loss of the association [41, 60]. The mechanism implemented in our model is overwriting the association, since the homeostatic mechanism drives the synaptic weights toward their initial value, thereby deleting the learned association with no chance of recovery, but only in the presence of olfactory input, eliciting MBON spikes. The resulting behavior during the extinction phase of the experiments presents itself in a similar way, while the underlying mechanism is different.

### Comparison of modeling results to experimental findings

A variety of experiments have demonstrated group-level acquisition curves that saturate over multiple training trials or with increasing duration of a single trial in olfactory conditioning [1, 51, 73–75]. To replicate larval behavior in reward learning experiments [1] with varying duration of the learning phase (fig. 5 A,B) we trained our model with an odor and reward in a paired vs. unpaired fashion (fig. 4 B). Real larvae show a strong odor preference even after a very short training and no significant increase in their preference when trained in a paired manner for longer periods of time [1, 67]. Instead, the animals trained in an unpaired protocol start out with a similarly high odor preference, which then decreases over time [1, 67]. This behavior is very counter-intuitive since the coincidence of odor and reward should yield an association of those two stimuli and thus an increased behaviorally expressed preference for the CS [8]. To resolve this contradiction, we include the observation that animals might not be naive to the training odor prior to the beginning of the experiments in the model. In that case, the animals would enter into the experiment with an already established reward prediction that would be violated during unpaired training. Three scenarios lend themselves as plausible causes of this effect: Firstly, accidental conditioning over the course of their lifespan during which they are raised on a food substrate while being exposed to air that carries many different odorants. Alternatively, or in fact, additionally, the animals might exhibit an innate preference for many odors [76–78]. Finally, the presence of the reward during reward-only phases might lead to an association of the experimental context with that reward (previously discussed by Saumweber et al. [67]). The resulting reward expectation (solely based on the always present context), unmet during the odor-only phases could lead to a prediction error signal. All three candidate explanations would yield a similar projection for the unpaired experimental protocol: A reward expectation acquired prior to the actual experiment would cause a violation of that expectation during odor-only trials of the unpaired experiments. In all three cases, the animal’s preferences might also generalize to a broader array of odors, leading to an overall preference for some odors, as observed experimentally. To test this hypothesis we pre-trained our model before simulating conditioning experiments (fig. 4 C) and observed that this allows us to reproduce the animal experiments (fig. 5 A,B). Including odor preference at the beginning of the experiment ensures the model not only behaves in accordance with the RW model [8], but also fits the animal experimental results [1]. A possible alternative explanation could be a sensory habituation process to the odor that might cause odor preferences to decrease over time, resulting in the observed patterns for unpaired learning. In the paired condition this effect might be abolished by the continued presentation of odor and reward together [79].

Thus far we have tested our model in experiments where the CS and US presentation were fully overlapping (paired conditions). We now consider different onset times, with the onset of the CS always preceding the onset of the US (fig. 3,C). For these experiments we used a shorter duration of 30*s* for both CS and US presentation, repeated over three acquisition trials to mimic experimental conditions in larval experiments [29, 66] that used optogenetic activation of DANs as a proxy for sugar reward. Similar to their experiments we show that the behavioral bias clearly depends on the temporal delay between CS and US (fig. 3,C). Complete temporal overlap of CS and US (ISI=0) does not seem to expend the full potential of learning the association, instead partial overlap yields stronger associations due to the extended window of opportunity for synaptic change triggered by the odor’s eligibility trace. In our model, the eligibility trace e(t) represents a molecular process that maintains the odor signal locally in the KC*>*MBON synapses (eqn. 2). Zeng et al. [80] demonstrate that feedback from the serotonergic dorsal paired medial neuron onto the KCs directly influences the length of the KC eligibility trace, making it a candidate mechanism for associative learning with a delayed US. Appetitive and aversive trace conditioning experiments have been conducted with larvae [29, 65, 66] and adult flies and other insects [74, 81–83]. In all of these experiments where the CS is presented before the US demonstrate that longer inter-stimulus intervals abolish learning of the CS-US association when no KC odor representation persists during the reinforcement period. In the cases of shorter intervals, the experimental data is not entirely conclusive. Either the odor preference was higher for partial or no overlap, compared with complete overlap [29, 83] or highest for complete overlap [51, 66, 74].

We also looked at the extent of reinforcement generalization to novel odors. Experiments have shown that associations between an odor and reinforcement generalize, to a varying extent, to other odors, as shown in experiments [63, 84]. Previous modeling experiments have also shown that reinforcement generalization depends on odor similarity in adult insects [48, 85–87]. In our larval model, we also demonstrate both generalization to other odors, as well as a loss in strength, compared to the training odor (fig. 3 A). We also show that the extent of the generalization depends on the similarity of the training and test odor, as measured by the overlap of the input patterns (fig. 1 D). The larval pathway with its relatively small coding space [53, 55] might be especially prone to such poor discriminative abilities.

### 0.1 Model predictions for behavioral experiments

Our approach targets two hypotheses: Firstly, symmetrical inhibitory and excitatory feedback from MBONs to DANs should yield a circuit capable of saturating learning curves as predicted by the RW model [8], due to PE [11] driving the learning process, which has also been suggested by previous models [20, 48–51]. Secondly, saturating learning curves, driven by PE should translate into (simulated) animal behavior, when comparing different training duration and intensities of reinforcement. We were able to test these hypotheses in model experiments, on the level of MB readout (behavioral bias, eqn. 4, fig. 2, 4)) and through the comparison of animal and simulated behavior of artificial larvae (fig. 5). While the simulation results fit nicely with the real larval behavior in an experiment with a varied training duration ([1], fig. 5), ultimately, the role of MBON*>*DAN feedback needs to be tested in behavioral animal experiments, directly manipulating this feedback. Some specific predictions that could be tested in such experiments are:

- Learning curves of individual animals should saturate over time when KC*>*MBON feedback is intact.
- When the MBON*>*DAN feedback is removed after some training, the learning curve should increase with a steeper slope and might not saturate.
- Increasing or decreasing the intensity of the odor or the reinforcement should lead to saturation on a higher or lower level, respectively.
- The removal of the KC*>*MBON feedback should weaken or abolish the saturation of the learning curve over time.

Based on our modeling results, we support the idea that the error computation between the prediction and reality of reinforcement is done in the DANs and relies on MBON*>*DAN feedback. Our hypotheses for experiments are based on this assumption. Nevertheless, some saturation, that is not based on PE, might still occur, even if MBON input to DANs is removed. The entire MB circuitry consists of many more elements than our model implementation and would presumably have additional mechanisms to ensure homeostatic balance and continued MBON input, potentially leading to some weaker form of saturation in the learning pro.

### Comparison with other MB models

Models of the learning in the MB, based on plasticity at the MB output synapses, without PE coding, have been around for some time, both for *Drosophila* [85, 87–89] and other insects [86, 90]. In all of these models, plasticity is mediated by the activity of modulatory neurons (e.g., dopaminergic), coinciding with either KC [86, 87] or coordinated KC and MBON activity [85, 88, 89]. These models can perform associative learning of a stimulus, paired with reinforcement [85–89], as well as more complicated forms of learning such as second order conditioning [89] and matching to sample [88] or reinforcement generalization tasks, the extent of which depends on the stimulus similarity [85, 86]. Additionally, some models were successfully tested in patterning tasks [85, 86], where combinations of stimuli are reinforced, while their individual components are not or vice versa. Models in which synaptic plasticity is driven not solely by the activity of modulatory neurons, but by a prediction error signal lend themselves to studying the evolution of learning over time (either over several trials, or in a continuous manner), and its dependency on the learning history. We hypothesize that such mechanisms for PE coding in the MB involve the modulatory DANs [19, 20, 32, 34, 41–44] and are based on MBON feedback to the DANs, serving as a manifestation of previous learning. Recently a number of modeling approaches have targeted the idea of PE coding in DANs in the adult *Drosophila* [48–51] as well as in the larval MB [20]. In these models, some form of MBON*>*DAN feedback is implemented, allowing these models to fulfill some of the predictions of the PE theory [8, 11]. One of the most fundamental predictions is the saturation of the learning curve across time, as the prediction error decreases, demonstrated in a trial-based manner in some of those models [48–51] as well as the loss of an acquired association [20, 48–50]. Some of the previously published models include mechanisms for either permanent loss of the association in memory or extinction (parallel associations in memory). Within the MB circuitry, the formation of a parallel extinction memory involves an additional DAN of opposite valence [20, 48–50], whereas complete loss is implemented as a process of changing the KC*>*MBON weights in the opposite direction of the learning process [51, 89], as done in our model. Additionally, some of these models capture temporal dynamics of learning experiments to some extent by utilizing eligibility traces in the KC*>*MBON synapses [20, 50, 51], to our understanding, none have tested these predictions in continuous experiments with spiking dynamics. Therefore, beyond the scope of these contributions, we implemented PE coding mechanistically in a fully spiking network equipped with synaptic eligibility that we train and test in continuous experiments to allow for the assessment of dynamic change in the model’s odor preference. In combination with a time-continuous behavioral simulation [52] during memory retention tests, this allowed for straightforward comparison with larval experiments.

Prediction error coding is not the only mechanism discussed in the literature to explain such phenomena in learning. Gkanias et al. tested a PE-based learning rule against a different dopamine-based learning rule that dos not require the presence of the CS as a reference point for expected reinforcement [87] in a more complex circuit model consisting of a number of interconnected micro-circuits. They show that both methods can produce a saturating learning curve across trials. Their alternative learning rule, embedded in a multi-compartment structure of the MB can also explain extinction, blocking, and second order conditioning, by relying on interactions between different MBONs and DANs that encode different memory processes.

### Outlook

Some experimentally observed effects in insect learning can not be captured by the RW model [8] and are thus not targeted by our model implementation. Among them are CS and US pre-exposure effects [91–94] that might be explained by changes either in attention to the CS or habituation to the CS or the US, caused by prolonged exposure prior to training, rather than changes in associative strength (for a review see [95]). Also interesting, but not directly predicted by the RW model [8] is the experimental observation of second order conditioning in adult *Drosophila* [96–99], where a second CS2 is paired with the CS, after this CS has acquired an association with the US. Through the CS2-CS pairing without the US, the CS2 acquires predictive power of the US. Different mechanisms have been proposed to be involved in causing this effect [98, 100]. Among them is an excitatory synaptic KC*>*DAN connection, strengthened during first order conditioning, that would allow the KC odor representation to activate the DAN as a substitute for reinforcement during the CS2-CS pairing. Exploring this phenomenon using network models could yield valuable insights into the *Drosophila* circuit, as well as aid in our general understanding of PE coding. Insect experiments have provided mixed evidence for other phenomena that can be predicted from the RW model, such as blocking [101–104] and hints at conditioned inhibition [105–107] that would be interesting to investigate. Furthermore, expanding the model to include different MB output compartments would offer a perspective to explore parallel associations regarding the same stimulus [41]. This could enable temporary loss of the learned association, while simultaneously retaining parallel memory for recovery (extinction vs. forgetting). Ultimately more possible directions arise from the major benefit of using a spiking model, which offers the potential to conduct experiments at high temporal resolution, instead of in a trial-based manner [20, 48–50]. In a future closed-loop approach that connects our continual learning MB model with the locomotory model in the full temporal resolution, we intend to simulate a behaving agent to investigate the temporal dynamics of adaptive behavior in analogy to the tracking experiments of real larva [73, 108–111].

## Methods

### Network model

All neurons are modeled as leaky integrate-and-fire neurons with conductance-based synapses. They elicit a spike, whenever the threshold *V_T_* is crossed(parameters provided in table S1). Each neuronal membrane potential *v_i_* is reset to the resting potential *V_r_* whenever a spike occurs, followed by an absolute refractory period of 2 ms, during which the neuron does not integrate any inputs. Any neuron from a given population (*v*^O^,*v*^P^,*v*^L^,*v*^K^,*v*^A^,*v*^M^,*v*^D^) is governed by the respective equation for ORNs, PNs, LNs, KCs, APL, MBONs and DANs (eqs. (5) to (11), fig. 1 A). Depending on the neuron type, in addition to a leak conductance *g_L_*, the equations consist of excitatory *g_e_* and inhibitory synaptic input *g_i_*. In the case of the DANs, one excitatory 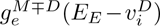 and inhibitory 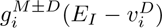 input represent the two types of MBON feedback for the reward and punishment encoding DAN, respectively. An additional spike-triggered adaptation conductance was implemented for ORNs, KCs, MBONs, and DANs (eqn. 12, [64]), in accordance with our current knowledge of the adaptive nature of ORNs in the larva [112] and the adult fly [113, 114]. Adaptation in KCs has so far only been demonstrated in other insects [115, 116]. In the model of these neurons, the adaptation conductance *g_Ia_* is increased with every spike and decays over time with *τ_Ia_*. The mechanism of synaptic plasticity is described in the results section (Learning through KC*>*MBON plasticity).

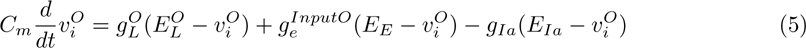

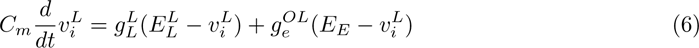

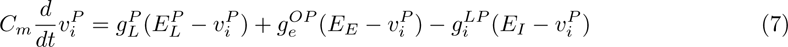

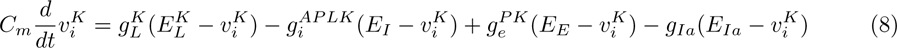

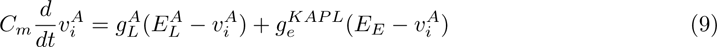

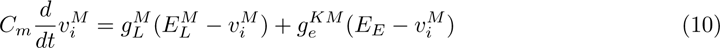

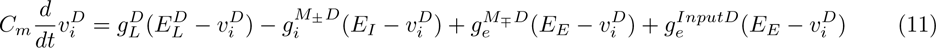

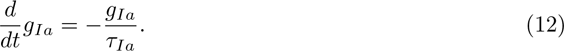

All code for the model implementation is accessible via https://github.com/nawrotlab/PEcodingDosophilaMB

We based our circuit model on the larval connectome both in terms of connectivity as well as numbers of neurons in each population [20, 53, 54] and introduced simplifications to support the mechanic investigation of the MBON*>*DAN feedback circuit and its role in PE coding and excluded a number of connections that have been demonstrated in the larva. Due to limited availability of anatomical, functional, and behavioral data most of our circuit implementation is based on the first instar larva [20, 53, 54], while the information on the APL connectivity within the circuit originates from studies on the third instar larva [29]. Behavioral experiments used for comparison with our simulation results were also performed with third instar larvae [1, 29, 66]. We demonstrate that our model based on the less developed circuit in the first instar larva is sufficient to reproduce animal behavior as observed in the older animals. From the anatomy of the first instar larva we excluded DAN*>*KC [54] and DAN*>*MBON synapses [54] that may play an additional role in learning-induced plasticity at KC*>*MBON synapses [54], the details of which are not fully known. Instead, we induce plasticity purely via the simulation of a neuromodulatory effect of the DANs onto the KC*>*MBON synapses ([54]). We also neglect recurrent interactions among KC themselves [54]. Many of these interactions affect KC that encode different sensory modalities, which are not included in our purely olfactory model. Furthermore, we simplified the connectivity between LNs and PNs [53] and between PNs and KCs to 2 *−* 6 PN inputs per KC, which excludes the set of KCs in the larva that receives exclusive input from only one PN [54]. This modification supported model robustness with respect to odor encoding within the small set of 72 KCs. Finally, from the population of *≈* 25 larval MBONs we only modeled two and correspondingly adapted KC*>*MBON synapses to provide both MBONs with input from all KCs.

### Sparse odor representation

We implemented four mechanisms supporting population- and temporal sparseness in the MB odor representation [64]. Population sparseness is defined as the activation of only a small subset of neurons by any given input [117]. In this circuit population sparseness is enhanced through lateral inhibition (via LNs), inhibitory APL feedback, and the divergent connectivity from PNs to a larger number of KCs [64]. Temporal sparseness indicates that an individual neuron responds with only a few spikes to a specific stimulus configuration [118–120], which supports encoding dynamic changes in the sensory environment [121, 122]. In our model temporal sparseness is facilitated by spike frequency adaptation, an adaptive process to prolonged stimulus exposure, in ORNs and KCs and by inhibitory feedback via the APL[64].

### Sensory input

In the olfactory pathway of larval *Drosophila* any odor activates up to *≈* 1/3 of ORNs, depending on its concentration [112, 123]. We implemented receptor input with stochastic point processes to ORNs via synapses to mimic the noise in a transduction process at the receptors. Each of the 21 receptor inputs is modeled according to a gamma process (shape parameter k= 3). The spontaneous firing rate of larval ORNs has been measured in the range of 0.2 *−* 7.9 Hz, depending strongly on odor and receptor type [123, 124]. ORNs in our model exhibit an average spontaneous firing rate of 8.92Hz (std=0.2). We constructed realistic olfactory input across the ORN population for amylacetate and 3-octanol by estimating ORN spike frequency from the calcium signals measured in the receptor neurons [112] (dilution of 10^-4^ [112]), ensuring the spike rates would not exceed the rates reported by [123]. They showed that using an even stronger odor concentration (dilution 10^-2^) ORN never exceeded a frequency of 200Hz. Due to the lower concentration used for amylacetate and 3-octanol (fig. 1 D) [112] in our experiments and because Kreher et al, 2005 measured only the first 0.5s after odor onset when the effects of spike frequency in ORNs are the weakest (leading to higher spike rates) we decided to use a maximum of 150Hz in odor activated ORNs. After generating the gamma process realizations we clipped multiple spikes occurring in each time step of the simulation discarding all but the first spike in each time step. Similar to the odor input, the presence of either reward or punishment in the experimental context was implemented as input to the DAN_+_/DAN_-_. Regular gamma spike trains (*k* = 10) were generated and clipped for the odor input.

To assess the effects of odor similarity on generalization we in addition created four artificial odors (A,B,C,D) (fig. 1 D) and quantified the pair-wise distances in ORN coding space using the cosine distance (eqn. 13), where vectors a and b each represent the input spike rate of two odors.

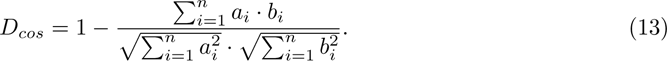

The cosine distance between odors A and B equals 0.21, 0.77 between odors A and C, and 0.99 between odors A and D. The comparison of amylacetate and 3-octanol yields a distance of 0.16.

### Experimental protocols

The experiments reported here belong to one of three categories. The first was performed to provide insight into the model and the effects of specific circuit functions on synaptic plasticity, and prediction error coding. To this end, we used amylacetate as the primary odor input. We varied the intensity of the reward via the frequency of gamma spike train, provided as input into the DAN_+_ (either 500Hz or 550Hz, resulting in an average output spike rate of 33.11/39.14Hz), and the learning rate *α*(0.6nS or 0.8nS). Additionally, MBON*>*DAN feedback was either enabled or disabled (fig. 1 A).

Experiments belonging to the second category were designed to replicate larva lab experiments to allow for a direct comparison with our model results. With these comparisons, we aim to validate the model and show to what extent our assumptions about the circuit functions allow us to recreate experimental data (fig. 5). Replicating lab experiments also provide more insights into the circuit mechanisms and offers alternative interpretations of the phenomena observed in data from animal experiments. Our implementations of the lab experiments were set up following the general procedure described in the Maggot Learning Manual [125]. Regardless of the specific protocol used in different experiments, larvae are placed into Petri dishes in groups of 30 animals. They are allowed to move around freely on the substrate that contains reinforcing substances, such as sugar or bitter tastants. During the entire time, they are subjected to specific odorants, emitted from two small containers in the dish to create permanent and uniformly distributed odor exposure within the dish. In the analogy of the experimental setting, in our simulated experiments, each model instance is trained individually through the concurrent presentation of olfactory stimulation and reward. One-minute intervals with only baseline ORN stimulation were included between training trials to simulate the time needed in the lab experiments for transferring larvae between Petri dishes. Unless otherwise specified and test phases refer to 3 min, during which only odors are presented. All simulations were implemented in the network simulator Brian2 [126].

### Realistic modeling of larval locomotion

Behavior during the testing phase of the olfactory learning experiment is simulated via the freely available python-based simulation platform Larvaworld (https://github.com/nawrotlab/larvaworld, [52]). A group of 30 virtual larvae is placed with random initial orientation around the center of a 100 mm diameter Petri dish and left to freely behave for 3 minutes. The previously conditioned odor is placed at one side of the dish, 10 mm from the arena’s boundary. Each larva features a bi-segmental body, supervised by a layered control architecture [52]. The basic layer of the control architecture is a locomotory model, capable of realistic autonomous exploration behavior. It consists of two coupled oscillators, one of which represents the crawling apparatus that generates forward velocity oscillations, resembling consecutive peristaltic strides [52]. The other oscillator generates alternating left and right lateral bending, manifested as oscillations of angular velocity [127]. The crawling and the bending oscillators are coupled via phase-locked suppression of lateral bending to capture the bend dependency on the stride-cycle phase during crawling (weathervaning). Finally, intermittent crawling is achieved by a superimposed intermittency module that generates alternating epochs of crawling and stationary pauses, with more headcasts for orientation during the latter [52]. Modulation of behavior due to sensory stimulation is introduced at the second, reactive layer of the control architecture. An odor signal can transiently alter both, the amplitude and frequency of the lateral bending oscillator, which biases free exploration towards approach or avoidance along an olfactory chemical gradient. This modulation of behavior is directly influenced via top-down signaling from the third, adaptive layer of the control architecture. In our approach, the spiking MB model populates the adaptive layer and its learning-dependent output, defined as the behavioral bias BB (i.e. the difference in MBON firing rates, eqn. 4), provides the top-down signal [36]. We formalize the gain of behavioral modulation as

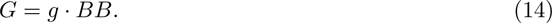

which is directly proportional to the behavioral bias and the additional proportionality factor *g* = 0.5.

A set of 10 *∗* 30 trained MB model instances is used to generate 10 groups of 30 simulated larvae. The preference index and the performance index [1] for these simulations are illustrated in figure 5. Preference indices (Pref) are computed individually for the paired and the unpaired experiments [1], based on the number of animals on each side (odor vs. empty) of the Petri dish at the end of the test phase.

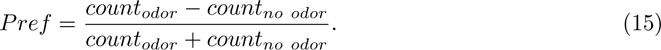

The Performance indices (PI) are computed from the preference indices of the paired and unpaired experiments [1].

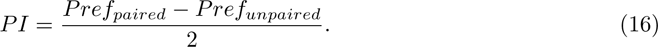

## Acknowledgements

This project is funded in parts by the German Research Foundation (DFG) within the Research Unit ‘Structure, Plasticity and Behavioral Function of the Drosophila mushroom body’ (DFG-FOR- 2705, grant no. 403329959, https://www.unigoettingen. de/en/601524.html to BG and MN) and by the Federal Ministry of Education and Research (BMBF, grant no. 01GQ2103A, ‘DrosoExpect’ to BG and MN). AMJ received additional support from the Research Training Group ‘Neural Circuit Analysis’ (DFG-RTG 1960, grant no. 233886668). We thank three anonymous reviewers for their valuable comments that helped us improve this manuscript.

---

**Figure S1:**
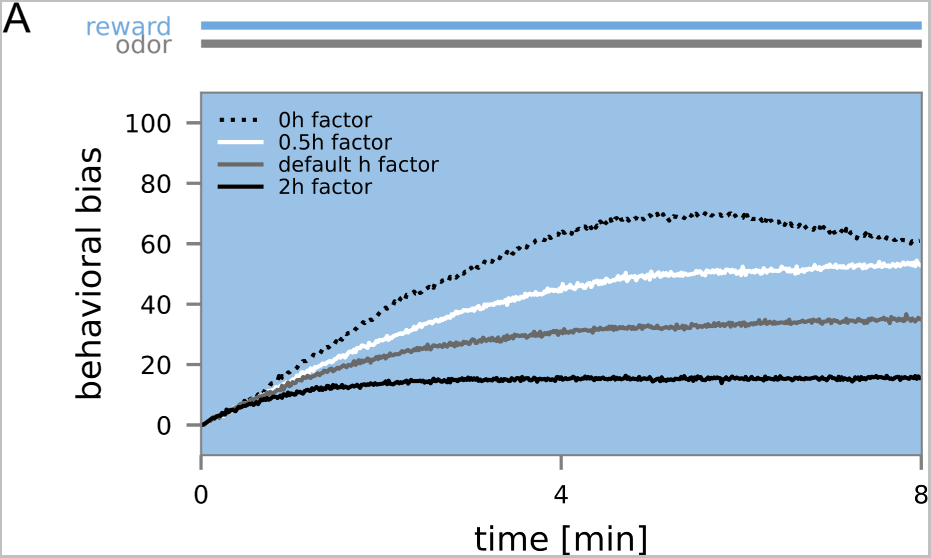
The effect of the homeostatic mechanism on the learning curve. (A) *N* = 30 model instances were trained with the odor amylacetate (CS) and reward (US, blue background). The odor preference (behavioral bias) was measured continuously in windows of 1 sec and averaged over all model instances. The learning rate was the same in all three experiments, while the magnitude of the homeostatic regulation h (eqn. 3,table S1) was either at its default value, at 0, or at half or twice the magnitude of the default value.

**Table S1:**
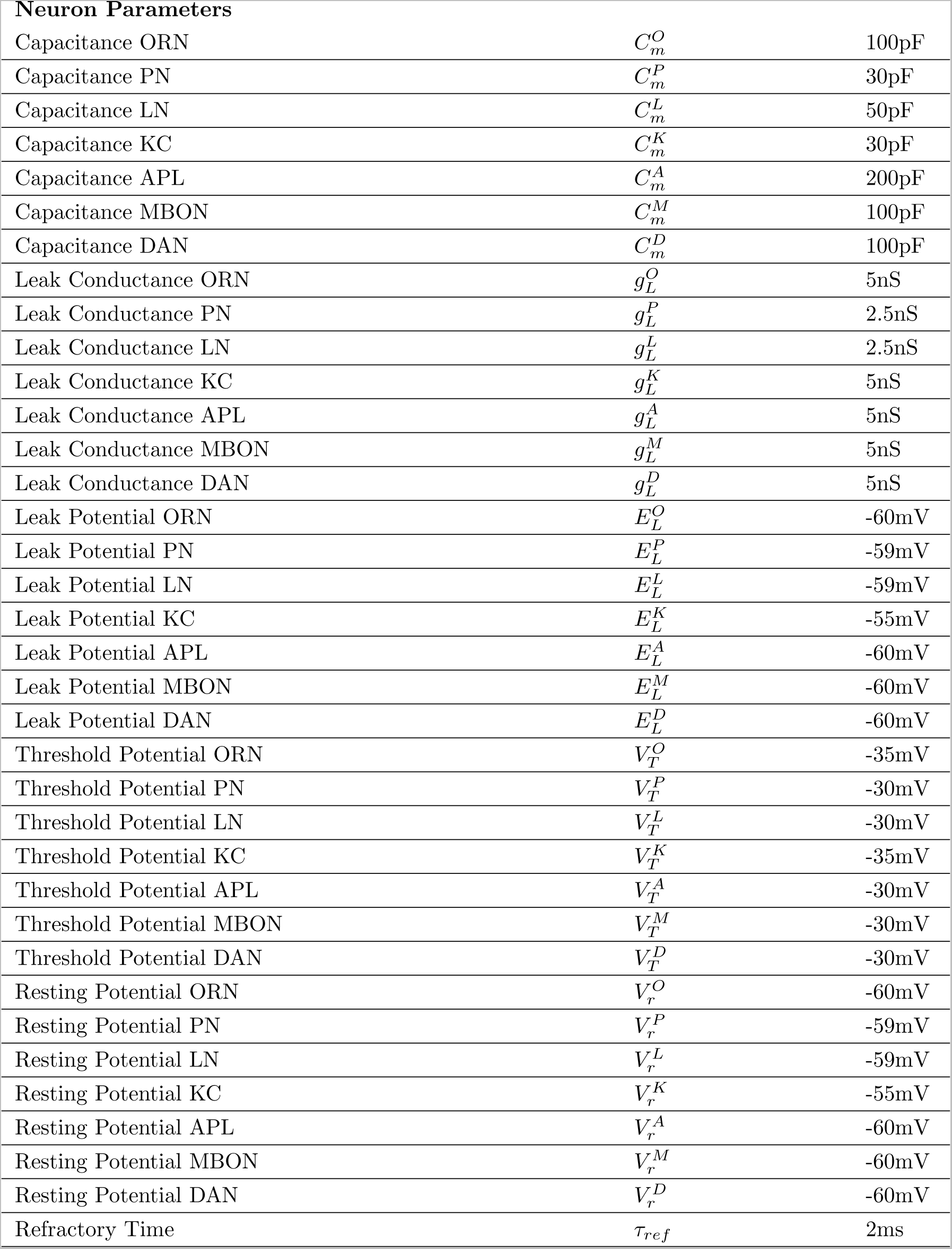
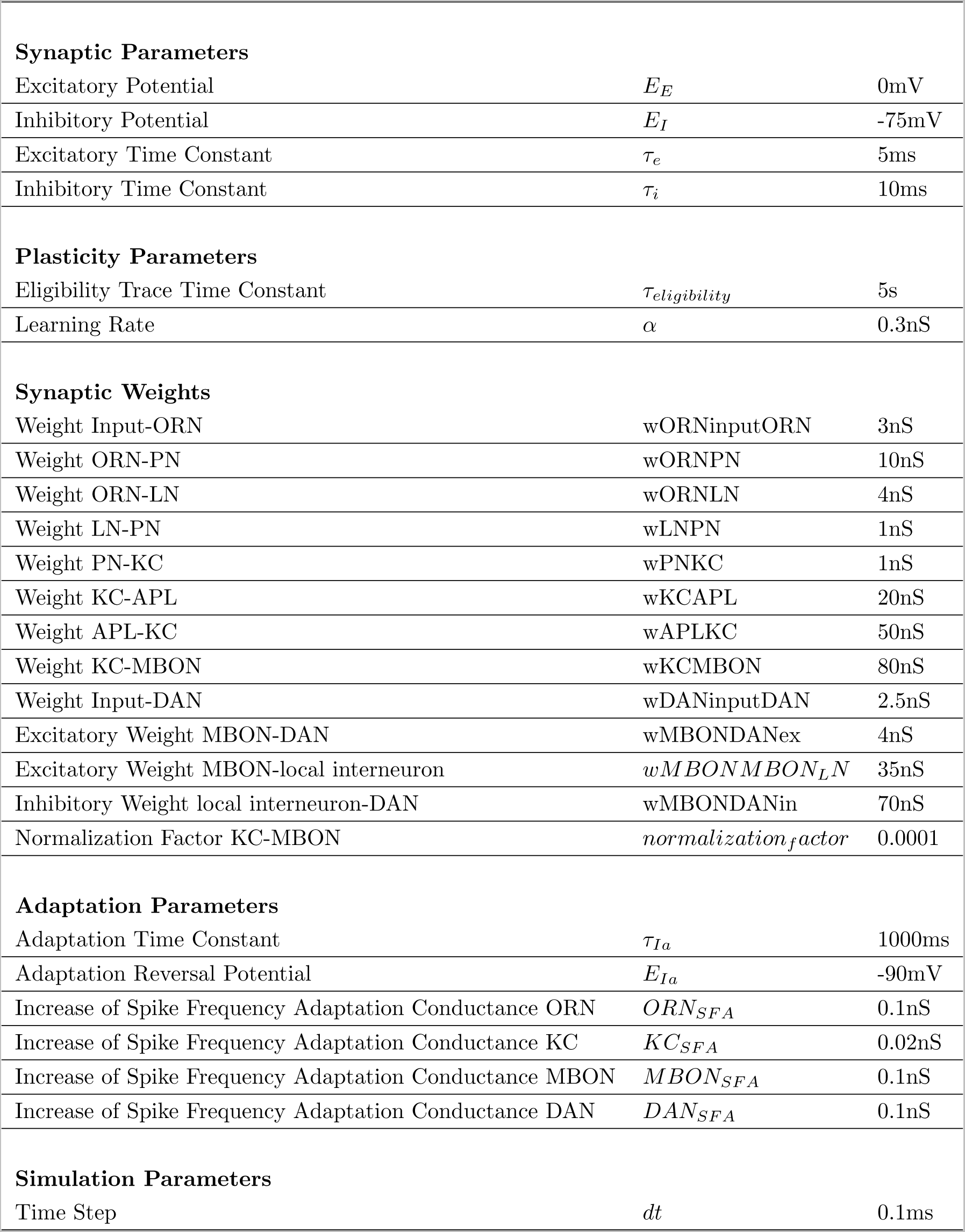
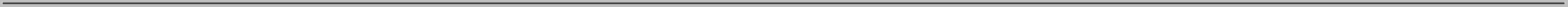
Network parameters

## Notes

### Competing Interest Statement

The authors have declared no competing interest.

### Summary of Updates

Updated version including reviewer feedback.

## References

[1] Aliće Weiglein, Florian Gerstner, Nino Mancini, Michael Schleyer, and Bertram Gerber. “One-trial learning in larval Drosophila”. In: Learning & Memory 26.4 (2019), pp. 109–120. doi: /10.1101/lm.049106.118.

[2] Martin Heisenberg. “Pattern recognition in insects”. In: Current Opinion in Neurobiology 5.4 (1995), pp. 475–481. issn: 09594388. doi: 10.1016/0959-4388(95)80008-5.

[3] Randolf Menzel. “The honeybee as a model for understanding the basis of cognition”. In: Nature Reviews Neuroscience 13.11 (2012), pp. 758–768. doi: 10.1038/nrn3357.

[4] Yoshinori Aso, Daisuke Hattori, Yang Yu, Rebecca M Johnston, Nirmala A Iyer, Teri-TB Ngo, Heather Dionne, LF Abbott, Richard Axel, Hiromu Tanimoto, et al. “The neuronal architecture of the mushroom body provides a logic for associative learning”. In: elife 3 (2014), e04577. doi: 10.7554/eLife.04577.

[5] Toshihide Hige, Yoshinori Aso, Mehrab N Modi, Gerald M Rubin, and Glenn C Turner. “Heterosynaptic plasticity underlies aversive olfactory learning in Drosophila”. In: Neuron 88.5 (2015), pp. 985–998. doi: /10.1016/j.neuron.2015.11.003.

[6] David Owald and Scott Waddell. “Olfactory learning skews mushroom body output pathways to steer behavioral choice in Drosophila”. In: Current opinion in neurobiology 35 (2015), pp. 178–184. doi: /10.1016/j.conb.2015.10.002.

[7] Martin F Strube-Bloss, Martin P Nawrot, and Randolf Menzel. “Mushroom body output neurons encode odor–reward associations”. In: Journal of Neuroscience 31.8 (2011), pp. 3129– 3140. doi: 10.1523/JNEUROSCI.2583-10.2011.

[8] Robert A Rescorla and Allan R Wagner. “A theory of Pavlovian conditioning : Variations in the effectiveness of reinforcement and non-reinforcement”. In: Classical Conditioning 2: Current Theory and Research. Ed. by William F. Black, Abraham H.; Prokasy. January 1972. New York, NY: Appelton-century-Crofts, 1972. Chap. 3, pp. 64–99.

[9] Peter D Balsam and Randy C Gallistel. “Temporal maps and informativeness in associative learning”. In: Trends in neurosciences 32.2 (2009), pp. 73–78. doi: 10.1016/j.tins.2008.10.004.

[10] Peter S Kaplan. “Importance of relative temporal parameters in trace autoshaping: From excitation to inhibition.” In: Journal of Experimental psychology: Animal behavior processes 10.2 (1984), p. 113.

[11] Leon Kamin. “Predictability, surprise, attention and conditioning”. In: Punsihment and aver-sive behavior. Ed. by Byron A. Campbell. New York: Appleton-Century-Crofts, 1969, 279–298.

[12] Wolfram Schultz. “Responses of midbrain dopamine neurons to behavioral trigger stimuli in the monkey”. In: Journal of neurophysiology 56.5 (1986), pp. 1439–1461. doi: 10.1152/jn.1986.56.5.1439.

[13] Genela Morris, David Arkadir, Alon Nevet, Eilon Vaadia, and Hagai Bergman. “Coincident but Distinct Messages of Midbrain Dopamine and Striatal Tonically Active Neurons”. In: Neuron 43.1 (2004), pp. 133–143. issn: 0896-6273. doi: 10.1016/J.NEURON.2004.06.012.

[14] Yoriko Takikawa, Reiko Kawagoe, and Okihide Hikosaka. “A possible role of midbrain dopamine neurons in short- and long-term adaptation of saccades to position-reward mapping”. In: Journal of Neurophysiology 92.4 (2004), pp. 2520–2529. doi: 10.1152/jn.00238.2004.

[15] Takemasa Satoh, Sadamu Nakai, Tatsuo Sato, and Minoru Kimura. “Correlated Coding of Motivation and Outcome of Decision by Dopamine Neurons”. In: Journal of Neuroscience 23.30 (2003), pp. 9913–9923. issn: 0270-6474. doi: 10.1523/JNEUROSCI.23-30-09913.2003.

[16] Jeremiah Y Cohen, Sebastian Haesler, Linh Vong, Bradford B Lowell, and Naoshige Uchida. “Neuron-type-specific signals for reward and punishment in the ventral tegmental area”. In: Nature 482.7383 (2012), pp. 85–88. doi: 10.1038/nature10754.

[17] Ariel Y Deutch, See-Ying Tam, and Robert H Roth. “Footshock and conditioned stress increase 3, 4-dihydroxyphenylacetic acid (DOPAC) in the ventral tegmental area but not substantia nigra”. In: Brain research 333.1 (1985), pp. 143–146. doi: 10.1016/0006-8993(85)90134-9.

[18] Scott Waddell. “Reinforcement signalling in Drosophila; dopamine does it all after all”. In: Current opinion in neurobiology 23.3 (2013), pp. 324–329. doi: 10.1016/j.conb.2013.01.005.

[19] Thomas Riemensperger, Thomas Völler, Patrick Stock, Erich Buchner, and André Fiala. “Punishment prediction by dopaminergic neurons in Drosophila”. In: Current Biology 15.21 (2005), pp. 1953–1960. issn: 09609822. doi: 10.1016/j.cub.2005.09.042.

[20] Claire Eschbach, Akira Fushiki, Michael Winding, Casey M Schneider-Mizell, Mei Shao, Rebecca Arruda, Katharina Eichler, Javier Valdes-Aleman, Tomoko Ohyama, Andreas S Thum, Bertram Gerber, Richard D Fetter, James W Truman, Ashok Litwin-Kumar, Albert Cardona, and Marta Zlatic. “Recurrent architecture for adaptive regulation of learning in the insect brain”. In: Nature Neuroscience 23.4 (2020), pp. 544–555. issn: 15461726. doi: 10.1038/s41593-020-0607-9.

[21] Martin Schwaerzel, Maria Monastirioti, Henrike Scholz, Florence Friggi-Grelin, Serge Birman, and Martin Heisenberg. “Dopamine and octopamine differentiate between aversive and appetitive olfactory memories in Drosophila”. In: Journal of Neuroscience 23.33 (2003), pp. 10495– 10502. doi: /10.1523/JNEUROSCI.23-33-10495.2003.

[22] Dale Corbett and Roy A. Wise. “Intracranial self-stimulation in relation to the ascending dopaminergic systems of the midbrain: A moveable electrode mapping study”. In: Brain Research 185.1 (1980), pp. 1–15. doi: 10.1016/0006-8993(80)90666-6.

[23] William R. Stauffer, Armin Lak, Aimei Yang, Melodie Borel, Ole Paulsen, Edward S. Boyden, and Wolfram Schultz. “Dopamine Neuron-Specific Optogenetic Stimulation in Rhesus Macaques”. In: Cell 166.6 (2016), 1564–1571.e6. doi: 10.1016/J.CELL.2016.08.024.

[24] Roy A Wise and P-P Rompre. “Brain dopamine and reward”. In: Annual review of psychology 40.1 (1989), pp. 191–225.

[25] Ilana B. Witten, Elizabeth E. Steinberg, Soo Yeun Lee, Thomas J. Davidson, Kelly A. Zalocusky, Matthew Brodsky, Ofer Yizhar, Saemi L. Cho, Shiaoching Gong, Charu Ramakrishnan, Garret D. Stuber, Kay M. Tye, Patricia H. Janak, and Karl Deisseroth. “Recombinase-driver rat lines: Tools, techniques, and optogenetic application to dopamine-mediated reinforcement”. In: Neuron 72.5 (2011), pp. 721–733. doi: 10.1016/j.neuron.2011.10.028.

[26] Chang Liu, Pierre Yves Plaaais, Nobuhiro Yamagata, Barret D. Pfeiffer, Yoshinori Aso, Anja B. Friedrich, Igor Siwanowicz, Gerald M. Rubin, Thomas Preat, and Hiromu Tanimoto. “A subset of dopamine neurons signals reward for odour memory in Drosophila”. In: Nature 2012 488:7412 488.7412 (2012), pp. 512–516. issn: 1476-4687. doi: 10.1038/nature11304.

[27] Adam Claridge-Chang, Robert D. Roorda, Eleftheria Vrontou, Lucas Sjulson, Haiyan Li, Jay Hirsh, and Gero Miesenböck. “Writing Memories with Light-Addressable Reinforcement Circuitry”. In: Cell 139.2 (2009), pp. 405–415. issn: 0092-8674. doi: 10.1016/J.CELL.2009.08.034.

[28] Christian König, Afshin Khalili, Mathangi Ganesan, Amrita P Nishu, Alejandra P Garza, Thomas Niewalda, Bertram Gerber, Yoshinori Aso, and Ayse Yarali. “Reinforcement signaling of punishment versus relief in fruit flies”. In: Learning & Memory 25.6 (2018), pp. 247–257. doi: 10.1101/lm.047308.118.

[29] Timo Saumweber, Astrid Rohwedder, Michael Schleyer, Katharina Eichler, Yi-Chun Chen, Yoshinori Aso, Albert Cardona, Claire Eschbach, Oliver Kobler, Anne Voigt, Archana Durairaja, Nino Mancini, Marta Zlatic, James W Truman, Andreas S Thum, and Bertram Gerber. “Functional architecture of reward learning in mushroom body extrinsic neurons of larval Drosophila”. In: Nature communications 9.1 (2018), p. 1104. doi: 10.1038/s41467-018-03130-1.

[30] Michael Schleyer, Aliće Weiglein, Juliane Thoener, Martin Strauch, Volker Hartenstein, Melisa Kantar Weigelt, Sarah Schuller, Timo Saumweber, Katharina Eichler, Astrid Rohwedder, et al. “Identification of dopaminergic neurons that can both establish associative memory and acutely terminate its behavioral expression”. In: Journal of Neuroscience 40.31 (2020), pp. 5990–6006. doi: 10.1523/JNEUROSCI.0290-20.2020.

[31] Christian Schroll, Thomas Riemensperger, Daniel Bucher, Julia Ehmer, Thomas Völler, Karen Erbguth, Bertram Gerber, Thomas Hendel, Georg Nagel, Erich Buchner, and André Fiala. “Light-induced activation of distinct modulatory neurons triggers appetitive or aversive learning in Drosophila larvae”. In: Current biology : CB 16 (17 2006), pp. 1741–1747. doi: 10.1016/j.cub.2006.07.023.

[32] Adithya E Rajagopalan, Ran Darshan, Karen L Hibbard, James E Fitzgerald, and Glenn C Turner. “Reward expectations direct learning and drive operant matching in Drosophila”. In: *bioRxiv* (2022), pp. 2022–05. doi: 10.1101/2022.05.24.493252.

[33] Clare E Hancock, Vahid Rostami, El Yazid Rachad, Stephan H Deimel, Martin P Nawrot, and André Fiala. “Visualization of learning-induced synaptic plasticity in output neurons of the Drosophila mushroom body *γ*-lobe”. In: Scientific Reports 12.1 (2022), p. 10421. doi: 10.1038/s41598-022-14413-5.

[34] Yoshinori Aso and Gerald M Rubin. “Dopaminergic neurons write and update memories with cell-type-specific rules”. In: Elife 5 (2016), e16135. doi: /10.7554/eLife.16135.

[35] Birgit Michels, Yi-chun Chen, Timo Saumweber, Dushyant Mishra, Hiromu Tanimoto, Benjamin Schmid, Olivia Engmann, and Bertram Gerber. “Cellular site and molecular mode of synapsin action in associative learning”. In: Learning & Memory 18.5 (2011), pp. 332–344. doi: 10.1101/lm.2101411.

[36] Claire Eschbach, Akira Fushiki, Michael Winding, Bruno Afonso, Ingrid V Andrade, Benjamin T Cocanougher, Katharina Eichler, Ruben Gepner, Guangwei Si, Javier Valdes-Aleman, et al. “Circuits for integrating learned and innate valences in the insect brain”. In: Elife 10 (2021), e62567. doi: /10.7554/eLife.62567.

[37] Wolfram Schultz, Paul Apicella, and Tomas Ljungberg. “Responses of monkey dopamine neurons to reward and conditioned stimuli during successive steps of learning a delayed response task”. In: Journal of neuroscience 13.3 (1993), pp. 900–913. doi: 10.1523/JNEUROSCI.13-03-00900.1993.

[38] Wolfram Schultz, Peter Dayan, and P Read Montague. “Getting formal with dopamine and reward”. In: Science 275.5306 (1997), pp. 1593–1599. doi: 10.1126/science.275.5306.1593.

[39] Kareem A Zaghloul, Justin A Blanco, Christoph T Weidemann, Kathryn McGill, Jurg L Jaggi, Gordon H Baltuch, and Michael J Kahana. “Human substantia nigra neurons encode unexpected financial rewards”. In: Science 323.5920 (2009), pp. 1496–1499. doi: 10.1126/science.1167342.

[40] Wei-Xing Pan, Robert Schmidt, Jeffery R Wickens, and Brian I Hyland. “Dopamine cells respond to predicted events during classical conditioning: evidence for eligibility traces in the reward-learning network”. In: Journal of Neuroscience 25.26 (2005), pp. 6235–6242. doi: /10.1523/JNEUROSCI.1478-05.2005.

[41] Johannes Felsenberg, Pedro F Jacob, Thomas Walker, Oliver Barnstedt, Amelia J Edmondson-Stait, Markus W Pleijzier, Nils Otto, Philipp Schlegel, Nadiya Sharifi, Emmanuel Perisse, et al. “Integration of parallel opposing memories underlies memory extinction”. In: Cell 175.3 (2018), pp. 709–722. doi: /10.1016/j.cell.2018.08.021.

[42] Makoto Mizunami, Kanta Terao, and Beatriz Alvarez. “Application of a prediction error theory to Pavlovian conditioning in an insect”. In: Frontiers in psychology 9 (2018), p. 1272. doi: /10.3389/fpsyg.2018.01272.

[43] Maria E. Villar, Miguel Pavão-Delgado, Marie Amigo, Pedro F. Jacob, Nesrine Merabet, Anthony Pinot, Sophie A. Perry, Scott Waddell, and Emmanuel Perisse. “Differential coding of absolute and relative aversive value in the Drosophila brain”. In: Current Biology 32.21 (2022), 4576–4592.e5. issn: 0960-9822. doi: 10.1016/j.cub.2022.08.058.

[44] Ilona C Grunwald Kadow and David Owald. “Decision making: Dopaminergic neurons for better or worse”. In: Current Biology 32.21 (2022), R1237–R1240. doi: 10.1016/j.cub.2022.09.043.

[45] Louis K Scheffer, C Shan Xu, Michal Januszewski, Zhiyuan Lu, Shin-ya Takemura, Kenneth J Hayworth, Gary B Huang, Kazunori Shinomiya, Jeremy Maitlin-Shepard, Stuart Berg, et al. “A connectome and analysis of the adult Drosophila central brain”. In: Elife 9 (2020), e57443. doi: /10.7554/eLife.57443.

[46] Nils Otto, Markus W Pleijzier, Isabel C Morgan, Amelia J Edmondson-Stait, Konrad J Heinz, Ildiko Stark, Georgia Dempsey, Masayoshi Ito, Ishaan Kapoor, Joseph Hsu, et al. “Input connectivity reveals additional heterogeneity of dopaminergic reinforcement in Drosophila”. In: Current Biology 30.16 (2020), pp. 3200–3211. doi: 10.1016/j.cub.2020.05.077.

[47] Michael Winding, Benjamin D Pedigo, Christopher L Barnes, Heather G Patsolic, Youngser Park, Tom Kazimiers, Akira Fushiki, Ingrid V Andrade, Avinash Khandelwal, Javier Valdes-Aleman, et al. “The connectome of an insect brain”. In: Science 379.6636 (2023), eadd9330. doi: 10.1126/science.add933.

[48] James E.M. Bennett, Andrew Philippides, and Thomas Nowotny. “Learning with reinforcement prediction errors in a model of the Drosophila mushroom body”. In: Nature Communications 12.1 (2021). issn: 20411723. doi: 10.1038/s41467-021-22592-4.

[49] Magdalena Springer and Martin P Nawrot. “A mechanistic model for reward prediction and extinction learning in the fruit fly”. In: Eneuro 8.3 (2021). doi: /10.1523/ENEURO.0549-20.2021.

[50] Linnie Jiang and Ashok Litwin-Kumar. “Models of heterogeneous dopamine signaling in an insect learning and memory center”. In: PLoS Computational Biology 17.8 (2021), e1009205. doi: /10.1371/journal.pcbi.1009205.

[51] Chang Zhao, Yves F Widmer, Sören Diegelmann, Mihai A Petrovici, Simon G Sprecher, and Walter Senn. “Predictive olfactory learning in Drosophila”. In: Scientific reports 11.1 (2021), pp. 1–17. doi: 10.1038/s41598-021-85841-y.

[52] Panagiotis Sakagiannis, Anna-Maria Jürgensen, and Martin P Nawrot. “A realistic locomotory model of Drosophila larva for behavioral simulations”. In: bioRxiv (2021). doi: 10.1101/2021.07.07.451470.

[53] Matthew E Berck, Avinash Khandelwal, Lindsey Claus, Luis Hernandez-Nunez, Guangwei Si, Christopher J Tabone, Feng Li, James W Truman, Rick D Fetter, Matthieu Louis, et al. “The wiring diagram of a glomerular olfactory system”. In: Elife 5 (2016). doi: /10.7554/eLife.14859.

[54] Katharina Eichler, Feng Li, Ashok Litwin-Kumar, Youngser Park, Ingrid Andrade, Casey M Schneider-Mizell, Timo Saumweber, Annina Huser, Claire Eschbach, Bertram Gerber, Richard D Fetter, James W Truman, Carey E Priebe, L F Abbott, Andreas S Thum, Marta Zlatic, and Albert Cardona. “The complete connectome of a learning and memory centre in an insect brain”. In: Nature 548.7666 (2017), pp. 175–182. issn: 1476-4687. doi: 10.1038/nature23455.

[55] Africa Couto, Mattias Alenius, and Barry J Dickson. “Molecular, anatomical, and functional organization of the Drosophila olfactory system”. In: Current Biology 15.17 (2005), pp. 1535– 1547. doi: /10.1016/j.cub.2005.07.034.

[56] Leslie B Vosshall and Reinhard F Stocker. “Molecular architecture of smell and taste in Drosophila”. In: Annual review of neuroscience 30 (2007), pp. 505–533. doi: 10.1146/annurev.neuro.30.051606.094306.

[57] Yoshinori Aso, Divya Sitaraman, Toshiharu Ichinose, Karla R Kaun, Katrin Vogt, Ghislain Belliart-Guérin, Pierre-Yves Plaçais, Alice A Robie, Nobuhiro Yamagata, Christopher Schnaitmann, et al. “Mushroom body output neurons encode valence and guide memory-based action selection in Drosophila”. In: Elife 3 (2014), e04580. doi: /10.7554/eLife.04580.

[58] Julien Séjourné, Pierre-Yves Plaçais, Yoshinori Aso, Igor Siwanowicz, Séverine Trannoy, Vladimiros Thoma, Stevanus R Tedjakumala, Gerald M Rubin, Paul Tchénio, KeiIto, et al. “Mushroom body efferent neurons responsible for aversive olfactory memory retrieval in Drosophila”. In: Nature neuroscience 14.7 (2011), pp. 903–910. doi: /10.1038/nn.2846.

[59] Clare E Hancock, Florian Bilz, and André Fiala. “In vivo optical calcium imaging of learning-induced synaptic plasticity in Drosophila melanogaster”. In: JoVE (Journal of Visualized Experiments) 152 (2019), e60288. doi: 10.3791/60288.

[60] Johannes Felsenberg, Oliver Barnstedt, Paola Cognigni, Suewei Lin, and Scott Waddell. “Reevaluation of learned information in Drosophila”. In: Nature 544.7649 (2017), pp. 240–244. doi: 10.1038/nature21716.

[61] Martin Schwaerzel, Martin Heisenberg, and Troy Zars. “Extinction antagonizes olfactory memory at the subcellular level”. In: Neuron 35.5 (2002), pp. 951–960. doi: 10.1016/S0896-6273(02)00832-2.

[62] Karyn M Myers and Michael Davis. “Behavioral and neural analysis of extinction”. In: Neuron 36.4 (2002), pp. 567–584. doi: 10.1016/S0896-6273(02)01064-4.

[63] Yi-chun Chen, Dushyant Mishra, Linda Schmitt, Michael Schmuker, and Bertram Gerber. “A behavioral odor similarity “space” in larval Drosophila”. In: Chemical senses 36.3 (2011), pp. 237–249. doi: 10.1093/chemse/bjq123.

[64] Anna-Maria Jürgensen, Afshin Khalili, Elisabetta Chicca, Giacomo Indiveri, and Martin P Nawrot. “A neuromorphic model of olfactory processing and sparse coding in the Drosophila larva brain”. In: Neuromorphic Computing and Engineering 1.2 (2021), p. 024008. doi: 10.1088/2634-4386/ac3ba6.

[65] Juliane Thoener, Aliće Weiglein, Bertram Gerber, and Michael Schleyer. “Optogenetically induced reward and ‘frustration’memory in larval Drosophila melanogaster”. In: Journal of Experimental Biology 225.16 (2022), jeb244565. doi: 10.1242/jeb.244565.

[66] Aliće Weiglein, Juliane Thoener, Irina Feldbruegge, Louisa Warzog, Nino Mancini, Michael Schleyer, and Bertram Gerber. “Aversive teaching signals from individual dopamine neurons in larval Drosophila show qualitative differences in their temporal “fingerprint””. In: Journal of Comparative Neurology 529.7 (2021), pp. 1553–1570. doi: /10.1002/cne.25037.

[67] Timo Saumweber, Jana Husse, and Bertram Gerber. “Innate attractiveness and associative learnability of odors can be dissociated in larval Drosophila”. In: Chemical senses 36.3 (2011), pp. 223–235. doi: /10.1093/chemse/bjq128.

[68] Michael Schleyer, Daisuke Miura, Teiichi Tanimura, and Bertram Gerber. “Learning the specific quality of taste reinforcement in larval Drosophila”. In: elife 4 (2015), e04711. doi: 10.7554/eLife.04711.

[69] Birgit Michels, Sören Diegelmann, Hiromu Tanimoto, Isabell Schwenkert, Erich Buchner, and Bertram Gerber. “A role for Synapsin in associative learning: the Drosophila larva as a study case”. In: Learning & Memory 12.3 (2005), pp. 224–231. doi: 10.1101/lm.92805.

[70] Mark E Bouton. “Context and behavioral processes in extinction”. In: Learning & memory 11.5 (2004), pp. 485–494. doi: 10.1101/lm.78804.

[71] Lingling Wang, Qi Yang, Binyan Lu, Lianzhang Wang, Yi Zhong, and Qian Li. “A behavioral paradigm to study the persistence of reward memory extinction in Drosophila”. In: Journal of genetics and genomics 46.12 (2019), pp. 599–601. doi: 10.1016/j.jgg.2019.11.001.

[72] Yukinori Hirano, Kunio Ihara, Tomoko Masuda, Takuya Yamamoto, Ikuko Iwata, Aya Takahashi, Hiroko Awata, Naosuke Nakamura, Mai Takakura, Yusuke Suzuki, et al. “Shifting transcriptional machinery is required for long-term memory maintenance and modification in Drosophila mushroom bodies”. In: Nature communications 7.1 (2016), p. 13471. doi: 10.1038/ncomms13471.

[73] Amanda Lesar, Javan Tahir, Jason Wolk, and Marc Gershow. “Switch-like and persistent memory formation in individual Drosophila larvae”. In: Elife 10 (2021), e70317. doi: /10.7554/eLife.70317.

[74] Tim Tully and William G Quinn. “Classical conditioning and retention in normal and mutant-Drosophila melanogaster”. In: Journal of Comparative Physiology A 157.2 (1985), pp. 263– 277.

[75] Kirsa Neuser, Jana Husse, Patrick Stock, and Bertram Gerber. “Appetitive olfactory learning in Drosophila larvae: Effects of repetition, reward strength, age, gender, assay type and memory span”. In: Animal Behaviour 69 (4 Apr. 2005), pp. 891–898. issn: 00033472. doi: 10.1016/j.anbehav.2004.06.013.

[76] Dennis Mathew, Carlotta Martelli, Elizabeth Kelley-Swift, Christopher Brusalis, Marc Gershow, Aravinthan DT Samuel, Thierry Emonet, and John R Carlson. “Functional diversity among sensory receptors in a Drosophila olfactory circuit”. In: Proceedings of the National Academy of Sciences 110.23 (2013), E2134–E2143. doi: 10.1073/pnas.130697611.

[77] Elane Fishilevich and Leslie B Vosshall. “Genetic and functional subdivision of the Drosophila antennal lobe”. In: Current Biology 15.17 (2005), pp. 1548–1553. doi: 10.1016/j.cub.2005.07. 066.

[78] Scott A Kreher, Dennis Mathew, Junhyong Kim, and John R Carlson. “Translation of sensory input into behavioral output via an olfactory system”. In: Neuron 59.1 (2008), pp. 110–124. doi: 10.1016/j.neuron.2008.06.010.

[79] Isabell Twick, John Anthony Lee, and Mani Ramaswami. “Olfactory habituation in Drosophila—odor encoding and its plasticity in the antennal lobe”. In: Progress in Brain Research 208 (2014), pp. 3–38. doi: 10.1016/B978-0-444-63350-7.00001-2.

[80] Jianzhi Zeng, Xuelin Li, Renzimo Zhang, Mingyue Lv, Yipan Wang, Ke Tan, Xiju Xia, Jinxia Wan, Miao Jing, Xiuning Zhang, et al. “Local 5-HT signaling bi-directionally regulates the coincidence time window for associative learning”. In: Neuron 111.7 (2023), pp. 1118–1135. doi: 10.1016/j.neuron.2022.12.034.

[81] Paul Szyszka, Christiane Demmler, Mariann Oemisch, Ludwig Sommer, Stephanie Biergans, Benjamin Birnbach, Ana F Silbering, and C Giovanni Galizia. “Mind the gap: olfactory trace conditioning in honeybees”. In: Journal of Neuroscience 31.20 (2011), pp. 7229–7239. doi: 10.1523/JNEUROSCI.6668-10.2011.

[82] Dana Shani Galili, Alja Lüdke, C Giovanni Galizia, Paul Szyszka, and Hiromu Tanimoto. “Olfactory trace conditioning in Drosophila”. In: Journal of Neuroscience 31.20 (2011), pp. 7240– 7248. doi: 10.1523/JNEUROSCI.6667-10.2011.

[83] Hiromu Tanimoto, Martin Heisenberg, and Bertram Gerber. “Event timing turns punishment to reward”. In: Nature 430.7003 (2004), pp. 983–983. doi: 10.1038/430983a.

[84] Dushyant Mishra, Matthieu Louis, and Bertram Gerber. “Adaptive adjustment of the generalization-discrimination balance in larval Drosophila”. In: Journal of neurogenetics 24.3 (2010), pp. 168–175. doi: /10.3109/01677063.2010.498066.

[85] Jan Wessnitzer, Joanna M Young, J Douglas Armstrong, and Barbara Webb. “A model of non-elemental olfactory learning in Drosophila”. In: Journal of computational neuroscience 32 (2012), pp. 197–212. doi: 10.1007/s10827-011-0348-6.

[86] Fei Peng and Lars Chittka. “A simple computational model of the bee mushroom body can explain seemingly complex forms of olfactory learning and memory”. In: Current Biology 27.2 (2017), pp. 224–230. doi: 10.1016/j.cub.2016.10.054.

[87] Evripidis Gkanias, Li Yan McCurdy, Michael N Nitabach, and Barbara Webb. “An incentive circuit for memory dynamics in the mushroom body of Drosophila melanogaster”. In: Elife 11 (2022), e75611. doi: 10.7554/eLife.75611.

[88] Paolo Arena, Luca Patané, Vincenzo Stornanti, Pietro Savio Termini, Bianca Zäpf, and Roland Strauss. “Modeling the insect mushroom bodies: Application to a delayed match-to-sample task”. In: Neural Networks 41 (2013), pp. 202–211. doi: 10.1016/j.neunet.2012.11.013.

[89] Faramarz Faghihi, Ahmed A Moustafa, Ralf Heinrich, and Florentin Wörgötter. “A computational model of conditioning inspired by Drosophila olfactory system”. In: Neural Networks 87 (2017), pp. 96–108. doi: 10.1016/j.neunet.2016.11.002.

[90] Ramón Huerta and Thomas Nowotny. “Fast and robust learning by reinforcement signals: Explorations in the insect brain”. In: Neural computation 21.8 (2009), pp. 2123–2151. issn: 0899-7667. doi: 10.1162/neco.2009.03-08-733.

[91] Andrew B Barron and Sarah A Corbet. “Pre-exposure affects the olfactory response of Drosophila melanogaster to menthol”. In: Entomologia experimentalis et applicata 90.2 (1999), pp. 175–181. doi: 10.1046/j.1570-7458.1999.00436.x.

[92] Vanesa M Fernández, Martin Giurfa, Jean-Marc Devaud, and Walter M Farina. “Latent inhibition in an insect: the role of aminergic signaling”. In: Learning & Memory 19.12 (2012), pp. 593–597. doi: /10.1101/lm.028167.112.

[93] Pedro F Jacob, Paola Vargas-Gutierrez, Zeynep Okray, Stefania Vietti-Michelina, Johannes Felsenberg, and Scott Waddell. “An opposing self-reinforced odor pre-exposure memory produces latent inhibition in Drosophila”. In: bioRxiv (2021). doi: 10.1101/2021.02.10.430636.

[94] Sathees BC Chandra, Jay S Hosler, and Brian H Smith. “Heritable variation for latent inhibition and its correlation with reversal learning in honeybees (Apis mellifera).” In: Journal of Comparative Psychology 114.1 (2000), p. 86. doi: 10.1037/0735-7036.114.1.86.

[95] RE Lubow, I Weiner, and Paul Schnur. “Conditioned attention theory”. In: Psychology of learning and motivation. Vol. 15. Elsevier, 1981, pp. 1–49. doi: 10.1146/annurev.neuro.30.051606.094306.

[96] Christopher J Tabone and J Steven de Belle. “Second-order conditioning in Drosophila”. In: Learning & Memory 18.4 (2011), pp. 250–253. doi: /10.1101/lm.2035411.

[97] Christian König, Afshin Khalili, Thomas Niewalda, Shiqiang Gao, and Bertram Gerber. “An optogenetic analogue of second-order reinforcement in Drosophila”. In: Biology Letters 15.7 (2019), p. 20190084. doi: 10.1101/lm.047308.118.

[98] Daichi Yamada, Daniel Bushey, Feng Li, Karen L Hibbard, Megan Sammons, Jan Funke, Ashok Litwin-Kumar, Toshihide Hige, and Yoshinori Aso. “Hierarchical architecture of dopaminergic circuits enables second-order conditioning in Drosophila”. In: Elife 12 (2023), e79042. doi: 10.7554/eLife.79042.

[99] Syed Abid Hussaini, Bernhard Komischke, Randolf Menzel, and Harald Lachnit. “Forward and backward second-order Pavlovian conditioning in honeybees”. In: Learning & Memory 14.10 (2007), pp. 678–683. doi: 10.1101/lm.471307.

[100] Isaac Cervantes-Sandoval, Anna Phan, Molee Chakraborty, and Ronald L Davis. “Reciprocal synapses between mushroom body and dopamine neurons form a positive feedback loop required for learning”. In: Elife 6 (2017), e23789. doi: /10.7554/eLife.23789.

[101] Kanta Terao and Makoto Mizunami. “Roles of dopamine neurons in mediating the prediction error in aversive learning in insects”. In: Scientific reports 7.1 (2017), pp. 1–9. doi: /10.1038/s41598-017-14473-y.

[102] Brian H Smith and Susan Cobey. “The olfactory memory of the honeybee Apis mellifera. II. Blocking between odorants in binary mixtures.” In: The Journal of experimental biology 195.1 (1994), pp. 91–108. doi: 10.1242/jeb.195.1.91.

[103] Robert S Thorn and Brian H Smith. “The olfactory memory of the honeybee Apis mellifera. III. Bilateral sensory input is necessary for induction and expression of olfactory blocking.” In: The Journal of experimental biology 200.14 (1997), pp. 2045–2055. doi: 10.1242/jeb.200. 14.2045.

[104] JS Hosler and Brian H Smith. “Blocking and the detection of odor components in blends”. In: Journal of Experimental Biology 203.18 (2000), pp. 2797–2806. doi: 10.1242/jeb.203.18.2797.

[105] K Takeda. “Classical conditioned response in the honey bee”. In: Journal of Insect Physiology 6.3 (1961), pp. 168–179.

[106] Kevin C Daly and Brian H Smith. “Associative olfactory learning in the moth Manduca sexta”. In: Journal of Experimental Biology 203.13 (2000), pp. 2025–2038. doi: 10.1242/jeb. 203.13.2025.

[107] Michael Schleyer, Markus Fendt, Sarah Schuller, and Bertram Gerber. “Associative learning of stimuli paired and unpaired with reinforcement: evaluating evidence from maggots, flies, bees, and rats”. In: Frontiers in psychology 9 (2018), p. 1494. doi: 10.3389/fpsyg.2018.01494.

[108] David Tadres and Matthieu Louis. “PiVR: An affordable and versatile closed-loop platform to study unrestrained sensorimotor behavior”. In: PLoS biology 18.7 (2020), e3000712. doi: /10.1371/journal.pbio.3000712.

[109] Benjamin Risse, Dimitri Berh, Nils Otto, Christian Klämbt, and Xiaoyi Jiang. “FIMTrack: An open source tracking and locomotion analysis software for small animals”. In: PLoS computational biology 13.5 (2017), e1005530. doi: /10.1371/journal.pcbi.1005530.

[110] Isabell Schumann and Tilman Triphan. “The pedtracker: An automatic staging approach for drosophila melanogaster larvae”. In: Frontiers in Behavioral Neuroscience (2020), p. 241. doi: /10.3389/fnbeh.2020.612313.

[111] Tomoko Ohyama, Tihana Jovanic, Gennady Denisov, Tam C Dang, Dominik Hoffmann, Rex A Kerr, and Marta Zlatic. “High-throughput analysis of stimulus-evoked behaviors in Drosophila larva reveals multiple modality-specific escape strategies”. In: PloS one 8.8 (2013), e71706. doi: /10.1021/acscatal.9b04293.

[112] Guangwei Si, Jessleen K. Kanwal, Yu Hu, Christopher J. Tabone, Jacob Baron, Matthew Berck, Gaetan Vignoud, and Aravinthan D.T. Samuel. “Structured Odorant Response Patterns across a Complete Olfactory Receptor Neuron Population”. In: Neuron 101.5 (2019), 950–962.e7. issn: 0896-6273. doi: 10.1016/J.NEURON.2018.12.030.

[113] Katherine I Nagel and Rachel I Wilson. “Biophysical mechanisms underlying olfactory receptor neuron dynamics”. In: Nature neuroscience 14.2 (2011), pp. 208–216. issn: 1546-1726. doi: 10.1038/nn.2725.

[114] Srinivas Gorur-Shandilya, Mahmut Demir, Junjiajia Long, Damon A Clark, and Thierry Emonet. “Olfactory receptor neurons use gain control and complementary kinetics to encode intermittent odorant stimuli”. In: Elife 6 (2017), e27670. doi: 10.7554/eLife.27670.

[115] Heike Demmer and Peter Kloppenburg. “Intrinsic Membrane Properties and Inhibitory Synaptic Input of Kenyon Cells as Mechanisms for Sparse Coding?” In: Journal of Neurophysiology 102.3 (2009), pp. 1538–1550. issn: 0022-3077. doi: 10.1152/jn.00183.2009.

[116] Jan Kropf and Wolfgang Rössler. “In-situ recording of ionic currents in projection neurons and Kenyon cells in the olfactory pathway of the honeybee”. In: PloS one 13.1 (2018), e0191425. doi: 10.1371/journal.pone.0191425.

[117] Bruno A. Olshausen and David J. Field. “Sparse coding of sensory inputs”. In: Current Opinion in Neurobiology 14.4 (2004), pp. 481–487. issn: 09594388. doi: 10.1016/j.conb.2004.07.007.

[118] Horace B Barlow. “Sensory mechanisms, the reduction of redundancy, and intelligence”. In: Mechanisation of thought processes. London: Her Majesty’s Stationery Office, 1959, pp. 535–539.

[119] Iori Ito, Rose Chik-Ying Ong, Baranidharan Raman, and Mark Stopfer. “Sparse odor representation and olfactory learning”. In: Nature neuroscience 11.10 (2008), pp. 1177–1184. issn: 1546-1726. doi: 10.1038/nn.2192.

[120] Roger Herikstad, Jonathan Baker, Jean Philippe Lachaux, Charles M. Gray, and Shih Cheng Yen. “Natural movies evoke spike trains with low spike time variability in cat primary visual cortex”. In: Journal of Neuroscience 31.44 (2011), pp. 15844–15860. issn: 02706474. doi: 10.1523/JNEUROSCI.5153-10.2011.

[121] Bilal Haider, Matthew R. Krause, Alvaro Duque, Yuguo Yu, Jonathan Touryan, James A. Mazer, and David A. McCormick. “Synaptic and Network Mechanisms of Sparse and Reliable Visual Cortical Activity during Nonclassical Receptive Field Stimulation”. In: Neuron 65.1 (2010), pp. 107–121. issn: 08966273. doi: 10.1016/j.neuron.2009.12.005.

[122] Chris Häusler, Alex Susemihl, and Martin Paul Nawrot. “Natural image sequences constrain dynamic receptive fields and imply a sparse code”. In: Brain research 1536 (2013), pp. 53–67. doi: /10.1016/j.brainres.2013.07.056.

[123] Scott A. Kreher, Jae Young Kwon, and John R. Carlson. “The molecular basis of odor coding in the Drosophila larva”. In: Neuron 46.3 (2005), pp. 445–456. issn: 08966273. doi: 10.1016/j.neuron.2005.04.007.

[124] Derek J Hoare, James Humble, Ding Jin, Niall Gilding, Rasmus Petersen, Matthew Cobb, and Catherine McCrohan. “Modeling peripheral olfactory coding in Drosophila larvae”. In: PLoS One 6.8 (2011), e22996. doi: /10.1371/journal.pone.0022996.

[125] Birgit Michels, Timo Saumweber, Roland Biernacki, Jeanette Thum, Rupert D.V. Glasgow, Michael Schleyer, Yi Chun Chen, Claire Eschbach, Reinhard F. Stocker, Naoko Toshima, Teiichi Tanimura, Matthieu Louis, Gonzalo Arias-Gil, Manuela Marescotti, Fabio Benfenati, and Bertram Gerber. “Pavlovian conditioning of larval Drosophila: An illustrated, multilingual, hands-on manual for odor-taste associative learning in maggots”. In: Front. Behav. Neurosci. 11.April (2017), pp. 1–6. issn: 16625153. doi: 10.3389/fnbeh.2017.00045.

[126] Marcel Stimberg, Romain Brette, and Dan FM Goodman. “Brian 2, an intuitive and efficient neural simulator”. In: Elife 8 (2019), e47314. doi: /10.7554/eLife.47314.

[127] Antoine Wystrach, Konstantinos Lagogiannis, and Barbara Webb. “Continuous lateral oscillations as a core mechanism for taxis in Drosophila larvae”. In: Elife 5 (2016). doi: 10.7554/elife.15504.

